# The long-term effects of chemotherapy on normal blood cells

**DOI:** 10.1101/2024.05.20.594942

**Authors:** Emily Mitchell, My H. Pham, Anna Clay, Rashesh Sanghvi, Sandra Pietsch, Joanne I. Hsu, Hyunchul Jung, Aditi Vedi, Sarah Moody, Jingwei Wang, Daniel Leonganmornlert, Michael Spencer Chapman, Nicholas Williams, Ellie Dunstone, Anna Santarsieri, Alex Cagan, Heather E. Machado, Joanna Baxter, George Follows, Daniel J Hodson, Ultan McDermott, Gary J. Doherty, Inigo Martincorena, Laura Humphreys, Krishnaa Mahbubani, Kourosh Saeb Parsy, Koichi Takahashi, Margaret A. Goodell, David Kent, Elisa Laurenti, Peter J. Campbell, Raheleh Rahbari, Jyoti Nangalia, Michael R. Stratton

## Abstract

In developed countries, ∼10% of individuals are exposed to systemic chemotherapy for cancer and other diseases. Many chemotherapeutic agents act by increasing DNA damage in cancer cells, triggering cell death. However, there is limited understanding of the extent and long-term consequences of collateral DNA damage to normal tissues. To investigate the impact of chemotherapy on mutation burdens and cell population structure of a normal tissue we sequenced blood cell genomes from 23 individuals, aged 3–80 years, treated with a range of chemotherapy regimens. Substantial additional mutation loads with characteristic mutational signatures were imposed by some chemotherapeutic agents, but there were differences in burden between different classes of agent, different agents of the same class and different blood cell types. Chemotherapy also induced premature changes in the cell population structure of normal blood, similar to those of normal ageing. The results constitute an initial survey of the long-term biological consequences of cytotoxic agents to which a substantial fraction of the population is exposed during the course of their disease management, raising mechanistic questions and highlighting opportunities for mitigation of adverse effects.

## Introduction

Over the course of a lifetime one in two people develop cancer. A longstanding approach to cancer treatment is systemic administration of a diverse group of cytotoxic chemicals, often collectively termed “chemotherapy”, which includes alkylating agents, platinum compounds, antimetabolites, topoisomerase inhibitors, vinca alkaloids and cytotoxic antibiotics^1^. Many are thought to exert their therapeutic effects by causing damage to DNA that, in turn, triggers death of malignant cells^2^. Approximately 30% of individuals with cancer, and thus approximately 10% of the whole population in developed countries, are exposed to chemotherapy at some point in their lifetime (Cancer Research UK, https://www.cancerresearchuk.org/health-professional/cancer-statistics-for-the-uk, accessed February 2024).

Chemotherapy can have long-term side effects on normal tissues. It confers an increased risk of cancers of blood^3–6^, lung, bladder and colon^7,8^ and is sometimes toxic to the kidney, blood, heart, brain, gastrointestinal tract, peripheral nervous system and gonads, engendering long-term deterioration in organ function^9–14^. There is limited understanding of the biological mechanisms underlying these sequelae of chemotherapy. However, it is plausible that some relate to consequences of DNA damage and thus could be elucidated through genome sequences of normal tissues, which may reveal changes in somatic mutation burdens or clonal composition, immediately or decades following chemotherapy.

Sequencing of cancers recurrent, or arising, after chemotherapy treatment has revealed variably elevated somatic mutation loads, in some instances characterised by distinctive mutational signatures^15–18^. However, there is little direct information concerning the mutagenic effects of chemotherapy on normal tissues *in vivo*. Studies of a small number of individuals show that normal colorectal epithelium, blood and sperm can show additional somatic mutation burdens after chemotherapy^19,20^. Furthermore, chemotherapy can alter the clonal structure of normal cell populations. This is illustrated in blood, where treatment increases the overall incidence of clonal haematopoiesis, favouring clones with driver mutations in the DNA damage response genes *PPM1D*, *TP53* and *CHEK2*^21–23^.

As part of a wider survey of the long-term impacts of chemotherapeutic agents on normal body tissues, here we investigate their effects on normal blood by whole genome sequencing of cells from chemotherapy-exposed individuals. Blood offers several desirable features in this regard, including the relative ease of randomly sampling cells from the whole tissue, the highly predictable mutation accumulation seen in unexposed individuals^24^, the opportunities to interrogate different cell subtypes and maturation states, and the feasibility of surveying changes in cell population clonal structure.

### Whole genome sequencing of chemotherapy-exposed blood

To conduct a primary survey of the landscape of chemotherapy effects on the genomes of normal blood cells we analysed 23 individuals who had collectively been exposed to multiple chemotherapy classes, multiple members of each class, at different ages and with variable time periods since exposure. These individuals, aged 3–80 years, had been treated with commonly used chemotherapy dosing regimens for haematological malignancies (Hodgkin lymphoma, n=2; follicular lymphoma, n=5; diffuse large B cell lymphoma, n=2; lymphoplasmacytic lymphoma, n=1; marginal zone lymphoma, n=1; multiple myeloma, n=1; acute myeloid leukaemia (AML), n=1) and solid cancers (colorectal carcinoma, n=9; neuroblastoma, n=1; lung cancer, n=1). One individual had been treated with chemotherapy for both multiple myeloma and colorectal carcinoma. The individual with AML had also been treated with chemotherapy for Behcet’s disease, a non-cancer condition (**Fig. 1a**; **Table S1**). Consistent with standard practice, most had received a combination of agents and, collectively, they had been exposed to 21 drugs from all of the main chemotherapy classes, including alkylating agents (cyclophosphamide, n=8; chlorambucil, n=2; bendamustine, n=5; procarbazine, n=2; melphalan, n=1), platinum agents (oxaliplatin, n=7; carboplatin, n=2; cisplatin, n=1), anti-metabolites (capecitabine, n=7; 5-fluorouracil, n=6; gemcitabine, n=1; cytarabine, n=1), topoisomerase-I inhibitors (irinotecan, n=5), topoisomerase-II inhibitors (etoposide, n=4; doxorubicin, n= 4; daunorubicin, n=1; mitoxantrone, n=1), vinca alkaloids (vincristine, n=7; vinblastine, n=1; vinorelbine, n=1), and cytotoxic antibiotics (bleomycin, n=1). Time periods from chemotherapy exposure to tissue sampling ranged from less than one month to six years for most cases. However, one individual sampled aged 48 years had been treated for Hodgkin lymphoma aged 10 and aged 47; and the individual sampled aged 43, post induction chemotherapy for AML, had also received long-term chlorambucil for Behcet’s disease diagnosed aged 13. In total 7 patients had received localised radiotherapy as part of their cancer therapy (**Table S1**). Results were compared with those from 9 healthy, chemotherapy non-exposed individuals (**Fig. 1a**; **Table S1**).

**Fig. 1:**
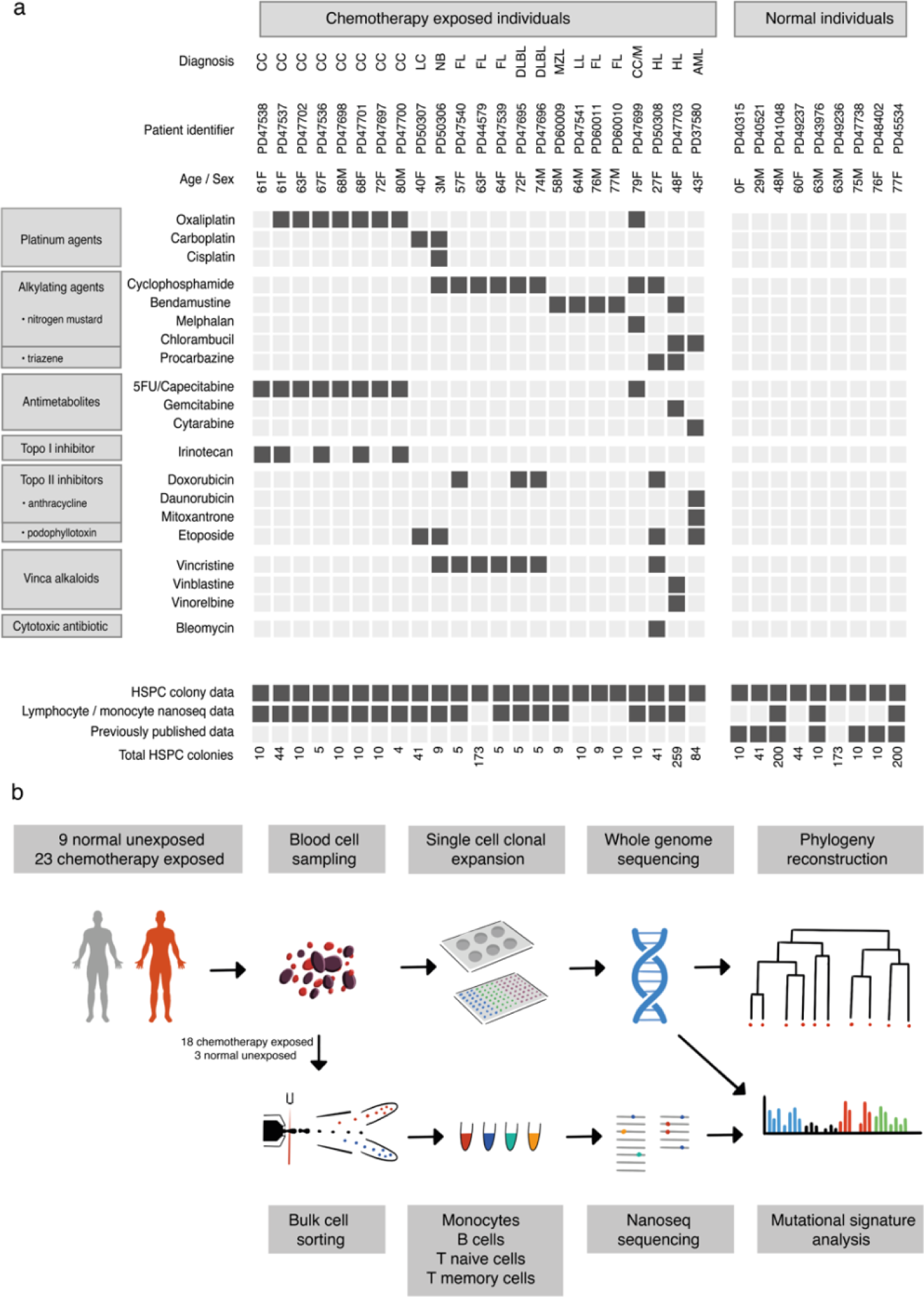
Donor information and experimental approach. **a**, Donor demographic details, chemotherapy exposure and sample information. CC = colorectal carcinoma, LC = lung cancer, NB = neuroblastoma, FL = follicular lymphoma, DLBL = diffuse large B cell lymphoma, MZL = marginal zone lymphoma, LL = lymphoplasmacytic lymphoma, M = multiple myeloma, HL = Hodgkin lymphoma, AML = acute myeloid leukaemia. **b**, Experimental approach.

Three experimental designs for detecting and analysing somatic mutations were used. First, 189 single cell-derived haematopoietic stem and progenitor cell (HSPC) colonies from the 23 chemotherapy-exposed individuals, and 90 from the nine controls, were expanded and individually whole genome sequenced at 23-fold average coverage to compare mutation burdens and mutational signatures. Second, from six individuals collectively exposed to a comprehensive range of the chemotherapeutic agents included in the study, a further 648 single cell colonies were whole genome sequenced (a total of 41–259 colonies each; mean sequencing depth 15-fold). These phylogenies were compared to similar sized phylogenies from five normal individuals across a similar age range (a total of 608 further colonies), to survey the effect of chemotherapy on the clonal structure of the HSPC population. Third, flow-sorted subpopulations of B cells, T memory cells, T naïve cells and monocytes from whole blood samples of 18 chemotherapy-exposed individuals and three unexposed normal individuals (**Fig. 1b**) were whole genome sequenced using duplex sequencing, which allows reliable identification of somatic mutations in polyclonal cell populations^25^. Somatic variant calling, mutational signature analysis and phylogenetic tree construction were performed as previously described (**Supplementary Methods**).

### Chemotherapy can increase haematopoietic cell somatic mutation burdens

Somatic single base substitution (SBS, also termed single nucleotide variant) mutations in HSPCs from normal adults accrue at a roughly constant rate of ∼18 per year, leading to a burden of ∼1500 SBSs in 80-year-old individuals^24^. HSPCs from 17/23 chemotherapy-exposed individuals showed elevated mutation burdens compared to those expected for their ages (**Fig. 2**). Four showed large increases of >1000 SBSs (**Fig. 2a**), 13 more modest increases of 200– 600 SBSs (**Fig. 2b**) and six no increases. The burdens of small indels in HSPCs were also increased in the four individuals with the greatest elevations in SBS burdens (**Extended Fig. 2a,b)**. Increases in structural variant and copy number changes were not observed, including in those individuals exposed to topoisomerase II inhibitors, which have been implicated in the development of secondary malignancies driven by specific oncogenic rearrangements^26^. (**Extended Fig. 2c,d**).

**Fig. 2:**
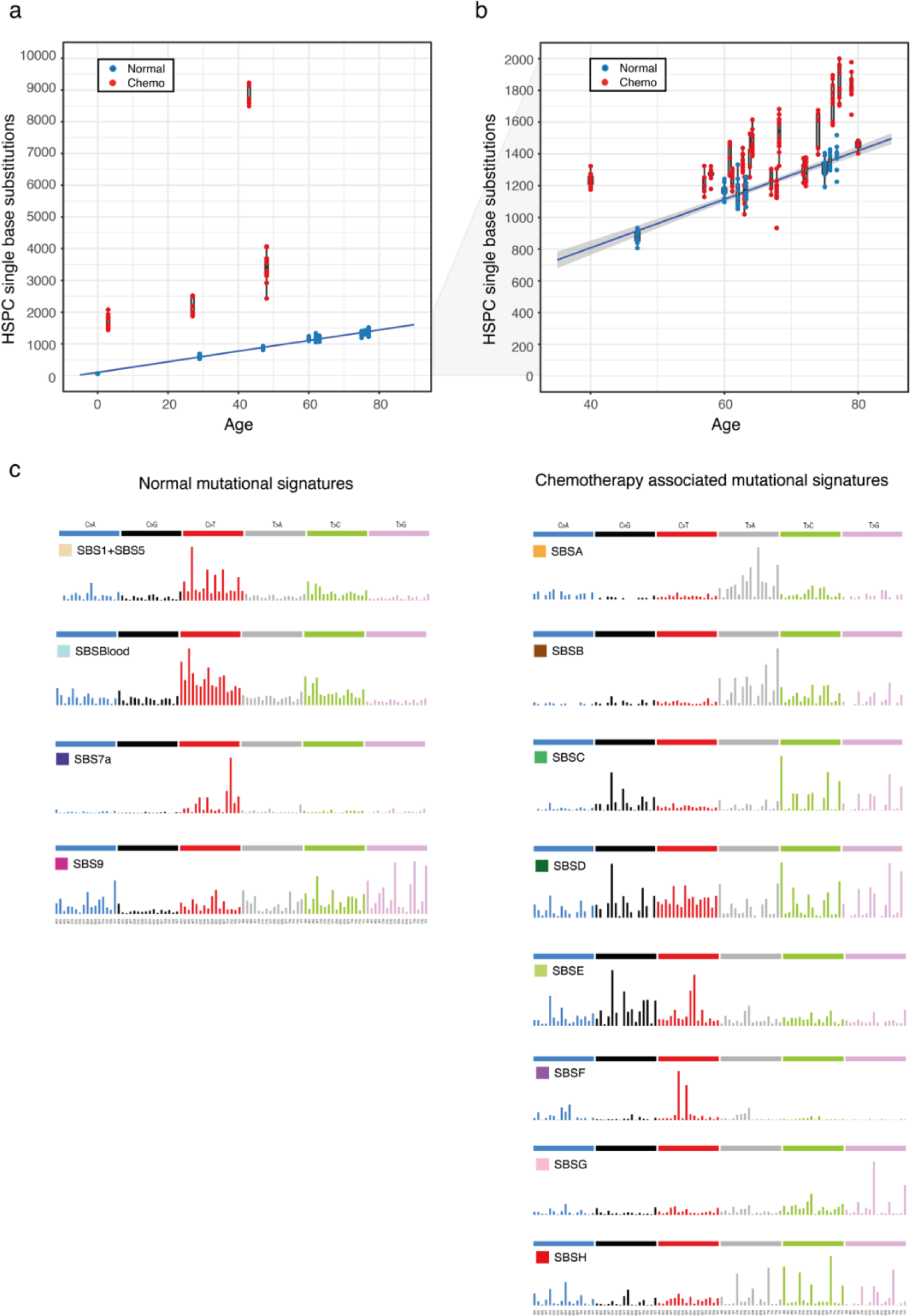
Mutational burden and mutational signatures in normal and chemotherapy exposed blood cells. **a**, Barplot of single base substitution burden with age (years) across normal (blue) and the four chemotherapy exposed (red) individuals with the highest indel burdens. The boxes indicate the median and interquartile range, the whiskers denote the minimum and maximum, with points representing outlying values. The blue line represents a regression of age on mutation burden across the unexposed individuals, with 95% CI shaded. **b**, Depiction of data as in a, but the y-axis is cut off at 2000 single base substitutions for better visualisation of the majority of the data. **c**, Mutational signatures extracted using HDP from the full dataset of normal and chemotherapy exposed HSPC colonies and duplex sequencing of bulk mature blood cell subsets.

Nineteen of the 23 chemotherapy-treated individuals received multiple agents. Therefore, it was in many cases uncertain which agents were responsible for the elevated mutation loads. To address this, we extracted mutational signatures from the SBS and indel mutation catalogues of chemotherapy-exposed individuals and controls, and estimated the contribution of each signature to the somatic mutations in the blood cells of each individual (**Fig. 2c; Extended Fig. 3**). We then used prior knowledge of previously described mutational signatures attributed to normal endogenous mutational processes, and to some mutagenic exposures^27^, as well as the specific chemotherapy regimens received by each individual in this study, to associate each signature with its putative causative agent.

Thirteen SBS mutational signatures were extracted. Five were similar to known signatures of normal HSPCs and mature lymphocytes: SBS1, characterised predominantly by C>T mutations at CG dinucleotides, and SBS5, which is relatively flat and featureless, are found in most normal cell types thus far studied; SBSBlood, a blood-specific signature predominant in HSPCs^28,29^; SBS7a, an ultraviolet light-caused signature found in memory T cells, which have presumably resided in skin during life^30^; and SBS9, a signature of somatic hypermutation found in B cells. Since two of the 13 signatures were predominantly composed of SBS1 and SBS5, these were combined for simplicity of depiction, generating a final total of 12 distinct signatures (**Fig. 2c**). Three indel mutational signatures were extracted. Two were similar to known indel signatures and present in both normal and chemotherapy-exposed individuals: the first comprised ID1 (ID1) and the second was a composite of ID3, ID5 and ID9 (ID3/5/9; **Extended Fig. 3)**.

Seven SBS mutational signatures were interpreted as being found exclusively in chemotherapy-treated individuals (**Fig. 2d)**, on the basis of accounting for <1% of mutations in HSPCs from adult controls (**Extended Fig. 4**). Four of these are novel, not represented in the COSMIC SBS mutational signature database. SBSA is likely due to the triazene alkylating agent procarbazine. There were three similar but distinct signatures relating predominantly to specific nitrogen mustard alkylating agents: SBSC to chlorambucil; SBSD to bendamustine; and SBSE to melphalan. SBSF is associated with the platinum agents cisplatin and carboplatin; and SBSG to the antimetabolite 5-fluorouracil or its prodrug capecitabine. The aetiologies of SBSB and SBSH are less clear cut and are discussed further below. Excess SBSs and specific SBS mutational signatures were not obviously associated with topoisomerase inhibitors (which cause DNA strand breaks), vinca-alkaloids (which inhibit microtubule formation during cell division) and the cytotoxic antibiotic bleomycin (which is thought to bind and cleave DNA). Only one high confidence indel mutational signature was found exclusively in chemotherapy-treated individuals: IDA, associated with procarbazine exposure.

SBSA contributed substantial additional mutation loads to blood cells from two individuals treated for Hodgkin lymphoma (PD50308 and PD44703) (**Fig. 3**). The only chemotherapy common to their treatment regimens was the alkylating agent procarbazine, no other individuals had been treated with procarbazine, and HSPC phylogenies indicated that SBSA mutations occurred early during PD44703’s life, consistent with procarbazine treatment aged 10 (**Extended Fig. 5a**). SBSA is similar to COSMIC signature SBS25 (cosine similarity 0.82), which has previously been associated with procarbazine^19,31^. An indel signature (IDA) was also identified as being most likely attributable to procarbazine, being found only in the two individuals treated with procarbazine; **Extended Fig. 3**. Alkylating agents cause alkyl DNA adducts, resulting in base mispairing and DNA breaks. Procarbazine is a triazene/hydrazine, monofunctional, alkylating agent.

**Figure 3:**
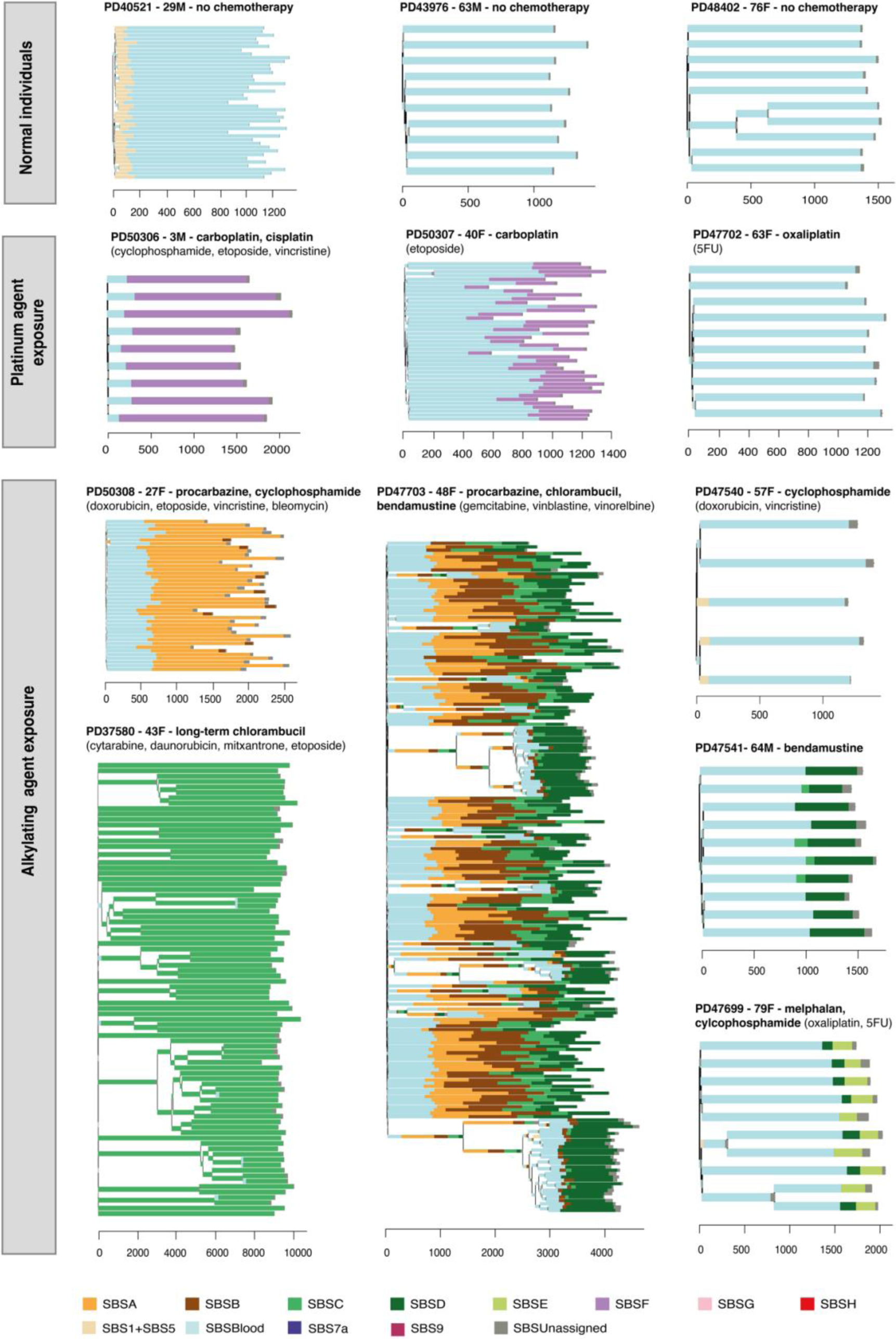
Phylogenetic trees and mutational signatures across a range of normal and chemotherapy exposed individuals. Phylogenies were constructed using shared mutation data and the algorithm MPBoot (Methods). Branch lengths correspond to SBS burdens. A stacked bar plot represents the signatures contributing to each branch with color code below the trees. SBSUnassigned indicates mutations that could not confidently be assigned to any reported signature.

SBSB is found predominantly in the individual exposed to chlorambucil, procarbazine and bendamustine (PD44703). Although SBSB, like SBSA (procarbazine), is predominantly composed of T>A substitutions, its cosine similarity to SBSA of 0.76 is not supportive of a strong similarity and the cosine similarities to SBSD (bendamustine; 0.68) and SBSC (chlorambucil; 0.62) indicate it is unlikely to be due to any of these in isolation. It is also present at low levels in the T memory cells of the other procarbazine exposed individual who was also exposed to cyclophosphamide (PD50308). Although its origin is unclear, it seems plausible that SBSB may result from an interaction between two classes of alkylating agent.

Of the nitrogen mustard associated signatures: SBSC contributed all mutations to the individual who received chlorambucil from childhood (PD37580); SBSD contributed all excess mutations to one of the individuals exposed only to bendamustine (PD60010) and was also present at much lower burden in a subset of cyclophosphamide-exposed individuals; SBSE was found only in the single individual exposed to low-dose melphalan (PD47699). Nitrogen mustard alkylating agents have two reactive sites and are, in consequence, bifunctional, forming intra- and inter-strand DNA cross-links in addition to simple adducts. The SBSC and SBSD signatures identified here are similar to a recently published mutational signature found in the germlines of two individuals whose fathers had been treated with two different nitrogen mustard agents (chlorambucil and iphosphamide) ^20,32^. SBSE is also similar (cosine similarity 0.84) to the previously described signature in multiple myeloma genomes with previous melphalan exposure^33–35^.

SBSF was found in individuals treated with carboplatin or cisplatin, and in a subset of oxaliplatin-treated individuals in whom it was present at much lower burdens. It is similar to COSMIC SBS31 (cosine similarity 0.96), which has previously been associated with prior platinum exposure in cancer genomes^32,36^ (**Fig. 3**). Platinum compounds act by binding DNA and forming intra- and inter-strand DNA crosslinks, in a similar manner to bifunctional alkylating agents. SBSF/SBS31 is, however, different from the bifunctional nitrogen mustard signatures, indicating that the patterns of DNA damage and/or DNA repair induced by platinum agents and nitrogen mustards differ.

SBSG is similar to COSMIC SBS17 (cosine similarity 0.91) which has previously been found in the genomes of cancers exposed to 5-fluorouracil^37^ and in the normal intestine of one 5-fluorouracil-exposed individual^19^. It was undetectable in HSPCs and found at highest burdens in lymphoid cells from individuals treated with 5-fluorouracil or its prodrug capecitabine (**Fig. 4**). 5-fluorouracil is a pyrimidine analogue mis-incorporated into DNA in place of thymine, consistent with causing a mutational signature characterised predominantly by mutations of thymine.

**Figure 4:**
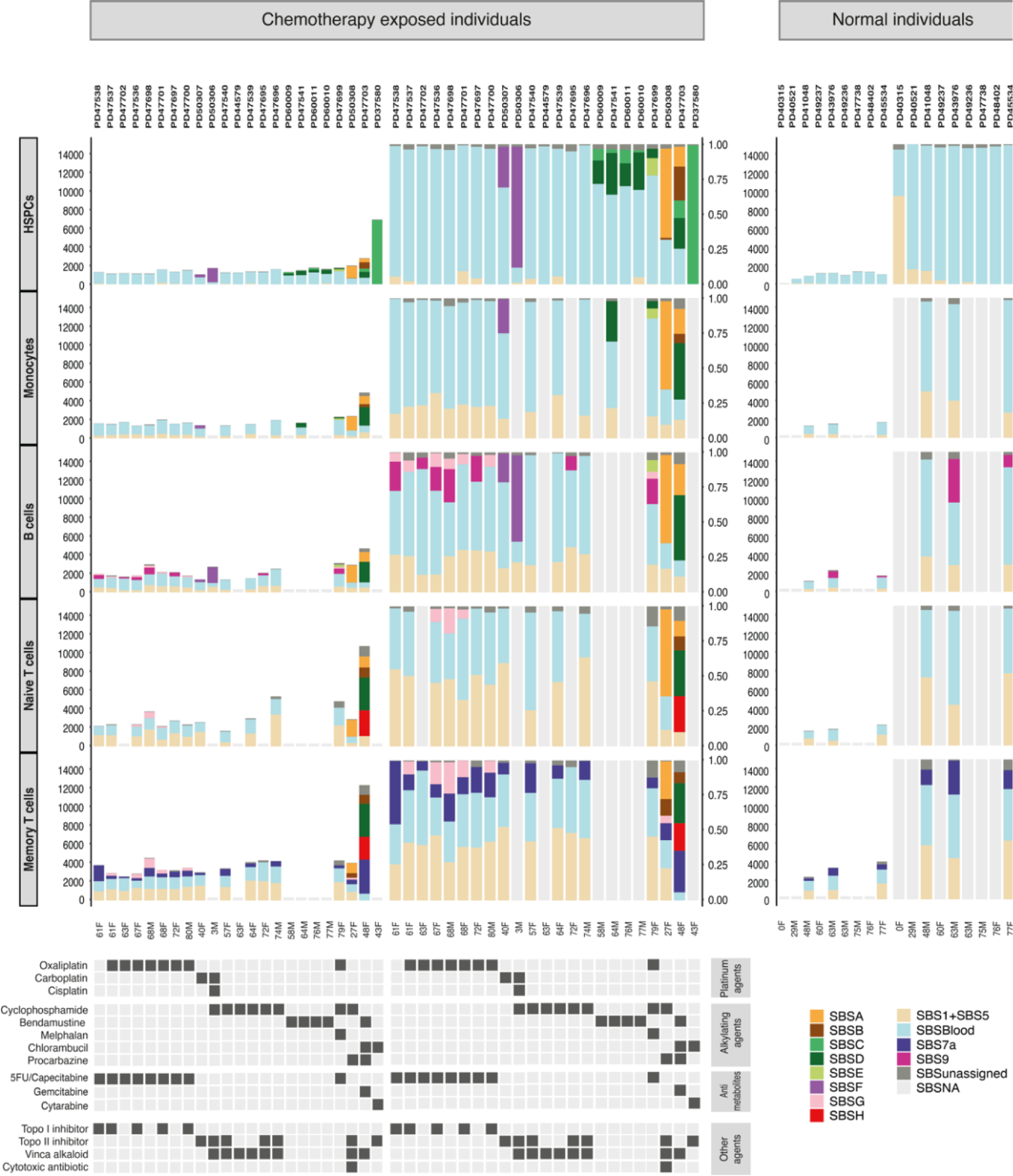
Mutation burden and SBS mutational signatures across different blood cell types. Stacked barplots represent the absolute contributions of each SBS mutational signature to the SBS mutation burden across cells types (left), compared to the proportionate contribution of each signature (right). HSPC data generated by pooling HSPC WGS colony data from each individual. Mature blood cell data generated using duplex sequencing of ∼40,000 cells of each type. For the normal unexposed individuals the T cell subset data is from CD4+ T cells whereas for the chemotherapy exposed individuals the T cell subsets contain both CD4+ and CD8+ cells. SBSUnassigned indicates mutations that could not confidently be assigned to any reported signature. SBSNA indicates duplex sequencing data is unavailable for this subset.

SBSH was detectable only in the T cells of a single individual who was also the only person to have received gemcitabine, a cytosine analogue. However, the origin of SBSH remains uncertain.

The isolation of multiple HSPC colonies from each individual allowed assessment of variation in mutagenic exposures across each of their HSPC populations. Overall, it was noteworthy how consistent the mutation burdens attributable to all the platinum agents, procarbazine and the various nitrogen mustards were across all sampled HSPCs from each individual (**Fig. 3**). This result indicates that, over the periods of chemotherapy exposure, there were few HSPCs continuously in niches or cell states protecting them from DNA damage and its consequences. The multiple HSPCs from each individual also allowed formation of phylogenetic trees allowing timing of mutagenic impacts. The phylogenetic timings were in keeping with the known periods of exposure; PD44703 with both early-life exposure to procarbazine and chlorambucil, and later-life exposure to bendamustine; PD37580 with both early and late life exposure to chlorambucil, and PD47699 with late-life exposure to melphalan (**Fig. 3**).

Although limited numbers of individuals, different drug combinations and different dose regimens preclude definitive evaluation, inclusion of individuals treated with different members of the same class of chemotherapy enabled preliminary comparison of their effects. Among nitrogen mustard alkylating agents, chlorambucil, bendamustine and melphalan caused substantially greater alkylating agent associated mutation burdens in normal blood cells than cyclophosphamide, which engendered only minimal (<5% and hence not shown in the figures) increases in mutation load (**Table S6,S7; Fig. 3; Extended Fig. 5b**). Similarly, carboplatin and cisplatin caused much higher SBSF mutation burdens than oxaliplatin, which conferred SBSF mutation burdens of <5% in all cases, despite prolonged oxaliplatin treatment (up to 22 cycles) in some individuals (**Table S6, S7; Fig. 3; Extended Fig. 6**). Therefore, chemotherapeutic agents of the same class, some used interchangeably in cancer treatment, may confer substantially different mutation burdens in normal blood cells.

Flow sorting of monocytes, B cells, T memory cells, and T naïve cells enabled investigation of the responses of different cell types to identical chemotherapy exposures. Overall, patterns of SBS signature burdens in monocytes were similar to HSPCs, whereas B and T lymphocytes showed differences for some agents. For example, SBSG, caused by 5-fluorouracil/capecitabine, contributed additional mutation burdens in B and T lymphocytes, but was undetectable in HSPCs and monocytes (**Fig. 4)**. In contrast, SBSF, caused by the platinum agents, contributed larger mutation burdens in HSPCs, monocytes and B cells than in T naïve and T memory cells, albeit we only have T cell data for one carboplatin exposed individual (**Fig. 4**). The mutation loads contributed by SBSA, caused by procarbazine, were similar across cell types. Therefore, some chemotherapeutic agents engender different mutation burdens in different cell types.

### Chemotherapy can change haematopoietic cell population structure

To investigate the effect of chemotherapies on the architecture of cell populations, extensive phylogenies of HSPCs from six chemotherapy-exposed individuals were generated and compared to non-exposed individuals of similar ages. An exemplar HSPC phylogeny of a normal, chemotherapy non-exposed 48-year-old shows only one barely-detectable clonal expansion and no ‘driver’ mutations in cancer genes (**Fig. 5a**). Such trees are typical of healthy middle-aged adults^24^.

**Figure 5:**
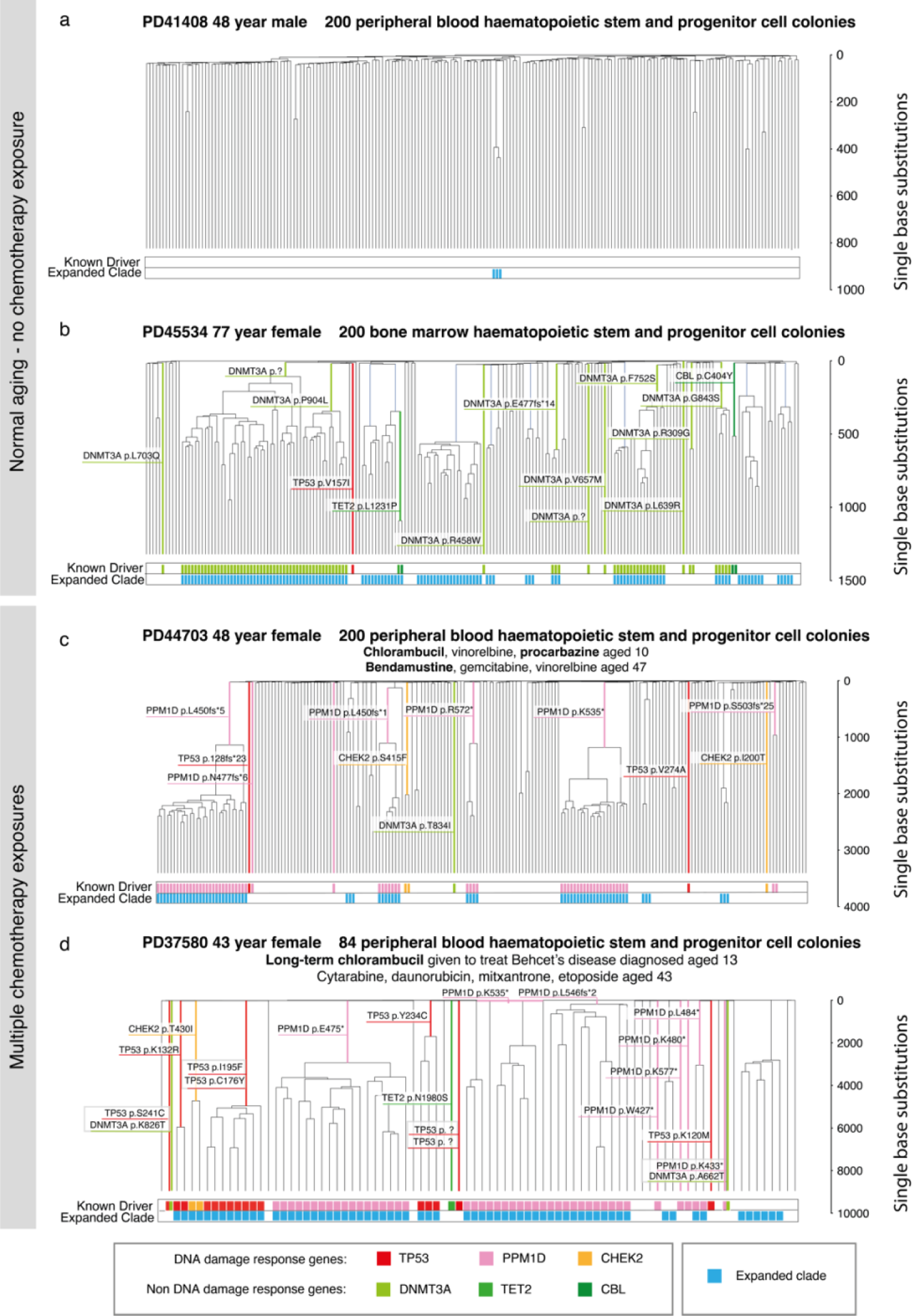
HSPC phylogenies for two normal unexposed and two chemotherapy exposed adult individuals. Phylogenies for two normal unexposed donors (top): one young and one elderly adult individual. Phylogenies for two young adult chemotherapy-treated individuals (bottom), who had both received more than one chemotherapy exposure. Phylogenies were constructed using shared mutation data and the algorithm MPBoot (Methods). Branch lengths reflect the number of mutations assigned to the branch with terminal branches adjusted for sequence coverage, and overall root-to-tip branch lengths have been normalized to the same total length (because all colonies were collected from a single time point). The y-axis represents the number of SBSs accumulating over time. Each tip on a phylogeny represents a single colony, with the respective numbers of colonies of each cell and tissue type recorded at the top. Onto these trees, we have layered clone and colony-specific phenotypic information. We have highlighted branches on which we have identified known oncogenic drivers in one of 18 clonal haematopoiesis genes (**Table S2**) color-coded by gene. A heat map at the bottom of each phylogeny highlights colonies from known driver clades coloured by gene, and the expanded clades (defined as those with a clonal fraction above 1%) in blue. **d**, The acute myeloid leukaemia derived from the bi-allelic *TP53* mutated clade carrying *TP53* p.I195F and *TP53* p.C176Y.

By contrast, a 48-year-old female (PD47703) treated for Hodgkin lymphoma with chlorambucil and procarbazine aged 10, and bendamustine aged 47, showed multiple independent clonal expansions carrying ‘driver’ mutations in the DNA damage response gene *PPM1D* (**Fig. 5c**). A similar pattern, with expanded *PPM1D* and *TP53* mutant clones, was observed in a 43-year-old female (PD37580) after long-term chlorambucil treatment (**Fig. 5d**). This pattern of multiple, large clonal expansions is characteristic of normal individuals aged >70 years^24^. However, in healthy elderly individuals, the clonal expansions exhibit predominantly *DNMT3A* and *TET2* driver mutations, or no apparent driver (**Fig. 5b**).

Chemotherapy could induce this prematurely-aged HSPC cell population profile by increasing mutation loads and/or by altering microenvironmental selection. Chemotherapy administered to elderly individuals favours survival of clonal haematopoiesis of indeterminate potential (CHIP) clones with driver mutations in *PPM1D*, *TP53* and *CHEK2*^21^, which usually predate the chemotherapy. Similarly, the HSPC phylogenetic tree of PD37580 indicates that at least two *PPM1D* driver mutations arose before chemotherapy was given during childhood (**Extended Fig. 7a**). Furthermore, in PD47703, comparison of two samples taken one year apart, during which additional chemotherapy (cyclophosphamide, doxorubicin and vincristine) had been administered, revealed a ∼50% increase in the size of pre-existing *PPM1D* mutated clones and no new mutant clones (**Extended Fig. 7b**). Thus, chemotherapy-induced changes in selection appear more influential than chemotherapy-induced creation of new driver mutations in generating the prematurely aged HSPC profile.

The prematurely aged architecture of the HSPC population was not observed in two other young adult individuals (PD50308 aged 29 and PD50307 aged 40) who received chemotherapy which caused substantial increases in mutation load and was administered two years or less before sampling (**Extended Fig. 8**), or in two further individuals, treated with cyclophosphamide and oxaliplatin (PD44579 aged 63 and PD47537 aged 61), who exhibited minimally increased mutation loads (**Extended Fig. 9**). It is, therefore, conceivable that multiple and/or prolonged chemotherapeutic exposures are required to generate the prematurely aged architecture. However, it is also possible that chemotherapy-engendered clonal expansions require decades to become detectable, as already demonstrated for clones under positive selection during normal ageing^24,38^.

The changes in clonal architecture resulting from chemotherapy exposure are significant for two reasons: firstly *PPM1D* mutant clones reduce the regenerative ability of the bone marrow in the setting of autologous bone marrow transplantation^39^, likely contributing to prolonged cytopenias and the death of PD47703 from infection 19 months post autograft. Secondly selection of *TP53* mutant clones confers a high risk of developing secondary myeloid malignancies, including AML as seen in PD37580, whose disease was treatment refractory and carried bi-allelic *TP53* mutations in association with a complex karyotype.

## Discussion

The results of this survey demonstrate that some commonly used chemotherapies, at dose regimens employed in clinical practice, increase somatic mutation burdens and alter the population structure of normal blood cells. Individuals with elevated mutation burdens have likely experienced very high mutation rates over short time periods. For example, an additional 1000 SBSs acquired in an HSPC due to chemotherapy administered during the course of one year is equivalent to a ∼50-fold increase in average mutation rate over the year, and it is plausible that mutation rates within hours or days of chemotherapy are even higher. The additional long-term mutation loads were also sometimes considerable. A three-year-old treated for neuroblastoma had >10-fold the number of somatic SBSs expected for his age, exceeding the burden in normal 80-year-olds.

The additional mutation burdens differed substantially both between chemotherapy classes and between agents of the same class. Since an important mechanism underlying the therapeutic effect of many chemotherapies is thought to be DNA damage induction, it is notable that different agents of the same class at their therapeutic doses have such different impacts on mutation generation in normal cells. The reasons for this are unclear, but may reflect subtle differences between agents in the nature of the DNA damage caused, the ability of damage to be repaired, the extent of induction of normal cell death and in the levels of normal cell exposure. For example, cyclophosphamide is thought to relatively spare HSPCs due to their higher levels of aldehyde dehydrogenase, an enzyme which inactivates a cyclophosphamide intermediary^40^. It may also be the case, however, that the extent of DNA damage does not directly correlate with the level of cytotoxicity of some chemotherapies.

The additional mutation burdens caused by chemotherapies are characterized by distinct mutational signatures, often shared by agents of the same class. The signatures are similar to those induced by the same agents in cancer cells, suggesting that the patterns of induced DNA damage, and its processing into mutations through DNA repair and replication, are similar in normal and cancer cells, even if tolerance of DNA damage by normal and cancer cells differs.

Increases in mutation loads imposed by chemotherapies differed between blood cell types and the profile of differences between cell types differed between chemotherapeutic agents. The mechanisms underlying these complex landscapes are uncertain, but may reflect intrinsic differences in the metabolic capabilities, DNA repair capacities and cell division rates of the different cell types.

Changes in the architecture of blood cell populations characterized by increasing dominance of large clones, often with driver mutations in cancer genes are a feature of normal ageing. Chemotherapy caused a similar pattern of change prematurely in some middle-aged adults, albeit with a different repertoire of mutated genes. However, these changes were not observed in all individuals. Whether these chemotherapy-induced changes in population architecture are contingent on long duration/multiplicity of treatment, or simply the passage of decades post treatment (which may allow clones with limited growth advantage under normal conditions to become detectable), requires further investigation.

In conclusion, some chemotherapies impose additional mutational loads and change the cell population structure of normal blood. Both impacts plausibly contribute to long-term consequences, including second malignancies, infertility, and loss of normal tissue resilience. Clinical data support this view, with the most mutagenic agents in this study having measurably greater long-term treatment toxicities. For example, of the bifunctional alkylating agents melphalan and chlorambucil are associated with higher risks of secondary malignancies than cyclophosphamide^3,41,42^. In addition, procarbazine has been associated with a particularly high risk of second cancer and infertility and is for this reason no longer used in the treatment of paediatric Hodgkin lymphoma^43^. Given that in many cancer types chemotherapeutic agents within a single class can be used interchangeably to achieve similar clinical outcomes^44–46^, it may be possible to prospectively utilise these types of data when improving existing regimens or developing new treatment protocols. In patients previously exposed to chemotherapy, knowledge of their altered mutational and clonal landscape could prompt discussions as to suitability for standard treatment protocols, particularly in the autologous transplant setting, and allow exploration of alternative options where appropriate.

The current study, together with others^17,32,47^, points to a future agenda for systematic genomic analysis of normal tissues after chemotherapy. This could incorporate multiple tissue sampling pre and post treatment, after short and long periods, across the range of chemotherapies, in substantial numbers of individuals, with detailed clinical and functional characterisation, and incorporating new sequencing technologies to enable feasibility at scale. A comprehensive prospective survey of this nature would improve understanding of the consequences of a widespread, self-administered mutagenic exposure in human populations and provide a scientific basis for optimising long-term patient health.

## Data availability

Additional data is available on github (https://github.com/emily-mitchell/chemotherapy/). Raw sequencing data is available on EGA (accession number WGS dataset EGAD00001015339 and Nanoseq dataset EGAD00001015340). The main data needed to reanalyse / reproduce the results presented is available on Mendeley Data (XXX)

## Code availability

Code is available on github: https://github.com/emily-mitchell/chemotherapy/

## Acknowledgements

This work was delivered as part of the Mutographs team supported by the Cancer Grand Challenges partnership funded by Cancer Research UK (*C98/A24032*). This work was supported by the Wellcome Trust grants 206194 and 220540/Z/20/A. D.J.H. was supported by a fellowship from Cancer Research UK (CRUK) (RCCFEL∖100072) and received core funding from Wellcome (203151/Z/16/Z) to the Wellcome-MRC Cambridge Stem Cell Institute and from the CRUK Cambridge Centre (A25117). This research was supported by the NIHR Cambridge Biomedical Research Centre (BRC-1215-555 20014). This research was jointly funded in whole or in part by the Wellcome Trust and the Royal Society. The views expressed are those of the authors and not necessarily those of the NIHR or the Department of Health and Social Care. E.L was supported by a Wellcome – Royal Society Sir Henry Dale Fellowship 107630/Z/15/Z (EL) and also by core support grants from Wellcome and the Medical Research Council (MRC) to the Wellcome-MRC Cambridge Stem Cell Institute 203151/Z/16/Z. Samples were provided by the Cambridge Blood and Stem Cell Biobank, which is supported by the Cambridge NIHR Biomedical Research Centre, Wellcome Trust - MRC Stem Cell Institute and the Cambridge Experimental Cancer Medicine Centre, UK. This research was supported by the Cambridge NIHR BRC Cell Phenotyping Hub. We are grateful to the donors, donor families and the Cambridge Biorepository for Translational Medicine for the gift of their tissue. The authors would like to thank Laura O’Neill, Kirsty Roberts, Katie Smith, Siobhan Austin-Guest and the staff of DNA Pipelines at the Wellcome Sanger Institute for their contribution. This research was funded in whole, or in part, by the Wellcome Trust. For the purpose of Open Access, the author has applied a CC BY public copyright licence to any Author Accepted Manuscript version arising from this submission. For the purpose of open access, the author has applied a CC-BY public copyright licence to any Author Accepted Manuscript version arising from this submission.

---

**Extended Data Fig. 1:**
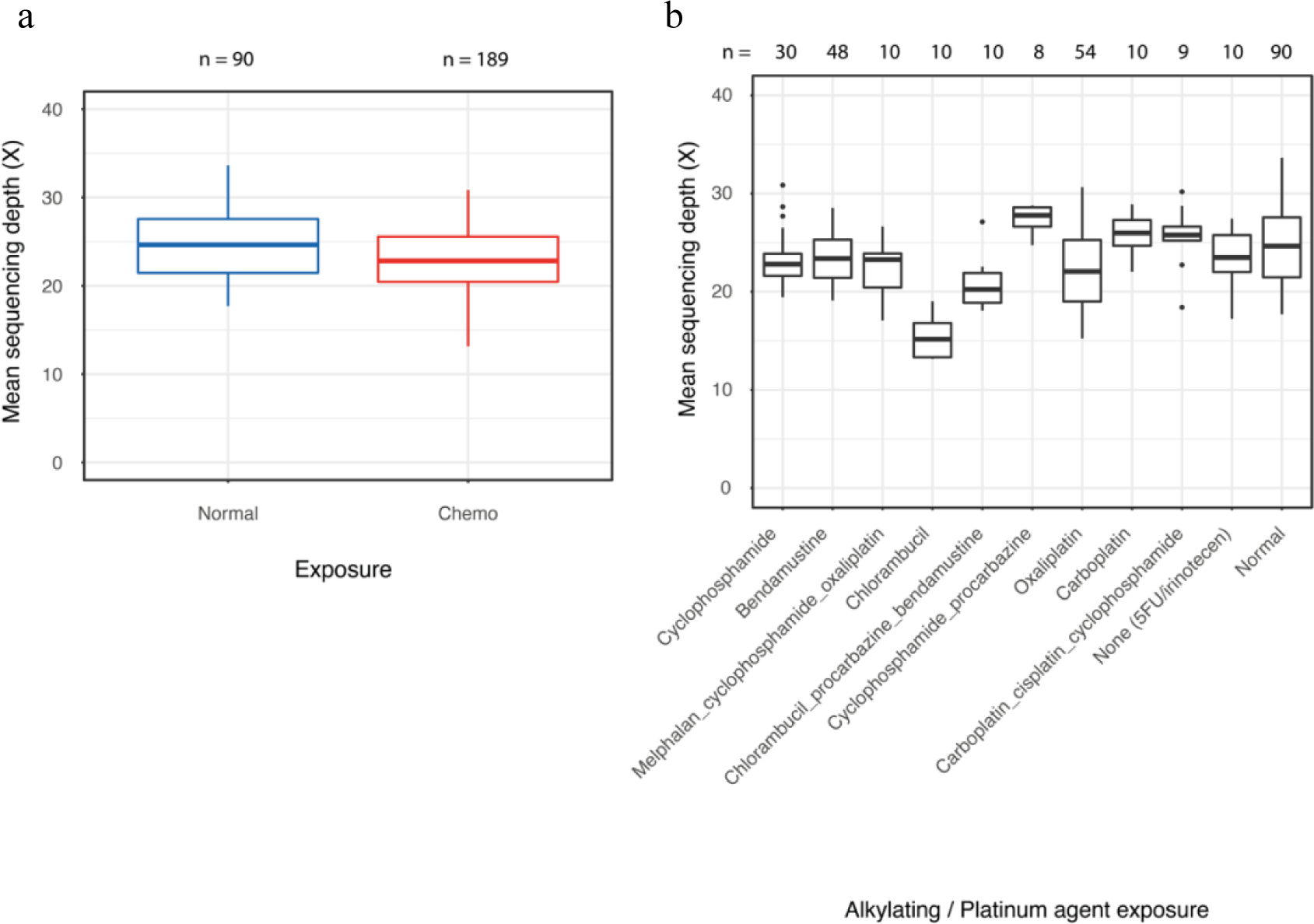
Mean sequencing depth in the normal and chemotherapy exposed HSPC colonies used for mutation burden analysis. a, Box plot representing the quartile distribution of mean sequencing depth in 90 colonies from normal (blue) and 189 colonies from chemotherapy exposed (red) individuals. b, Boxplot comparing the mean sequencing depth between Alkylating/ Platinum agent exposed and non-exposed colonies. The number of colonies in each agent group are shown at the top of the plot.

**Extended Data Fig. 2:**
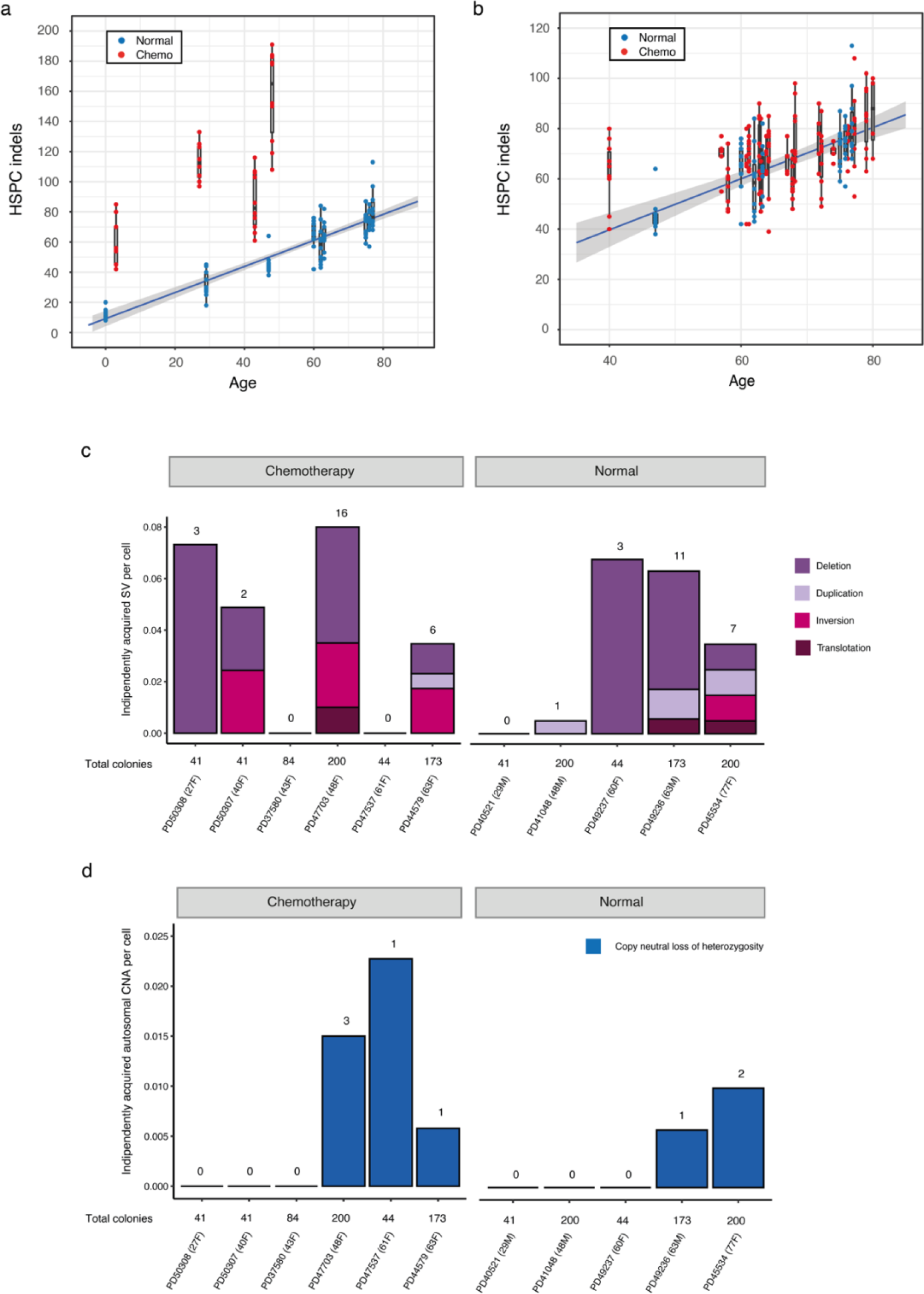
Indel mutational burden in normal and chemotherapy exposed HSPCs. a, Barplot of small indel burden with age (years) across normal (blue) and the four chemotherapy exposed (red) individuals with the highest indel burdens. The boxes indicate the median and interquartile range, the whiskers denote the minimum and maximum, with points representing outlying values. The blue line represents a regression of age on mutation burden, with 95% CI shaded. b, Depiction of data as in a, but the y-axis is cut off at 120 indels for better visualisation of the majority of the data. c,d, Bar plots showing the number of structure variant types (c) and the number of the number of independently acquired autosomal copy number aberrations (CNAs) (d) in each individual from chemotherapy and normal groups. The absolute number of events found in each individual is shown at the top of each bar. Individuals and the total number of isolated colonies are sorted by age within each group.

**Extended Data Fig. 3:**
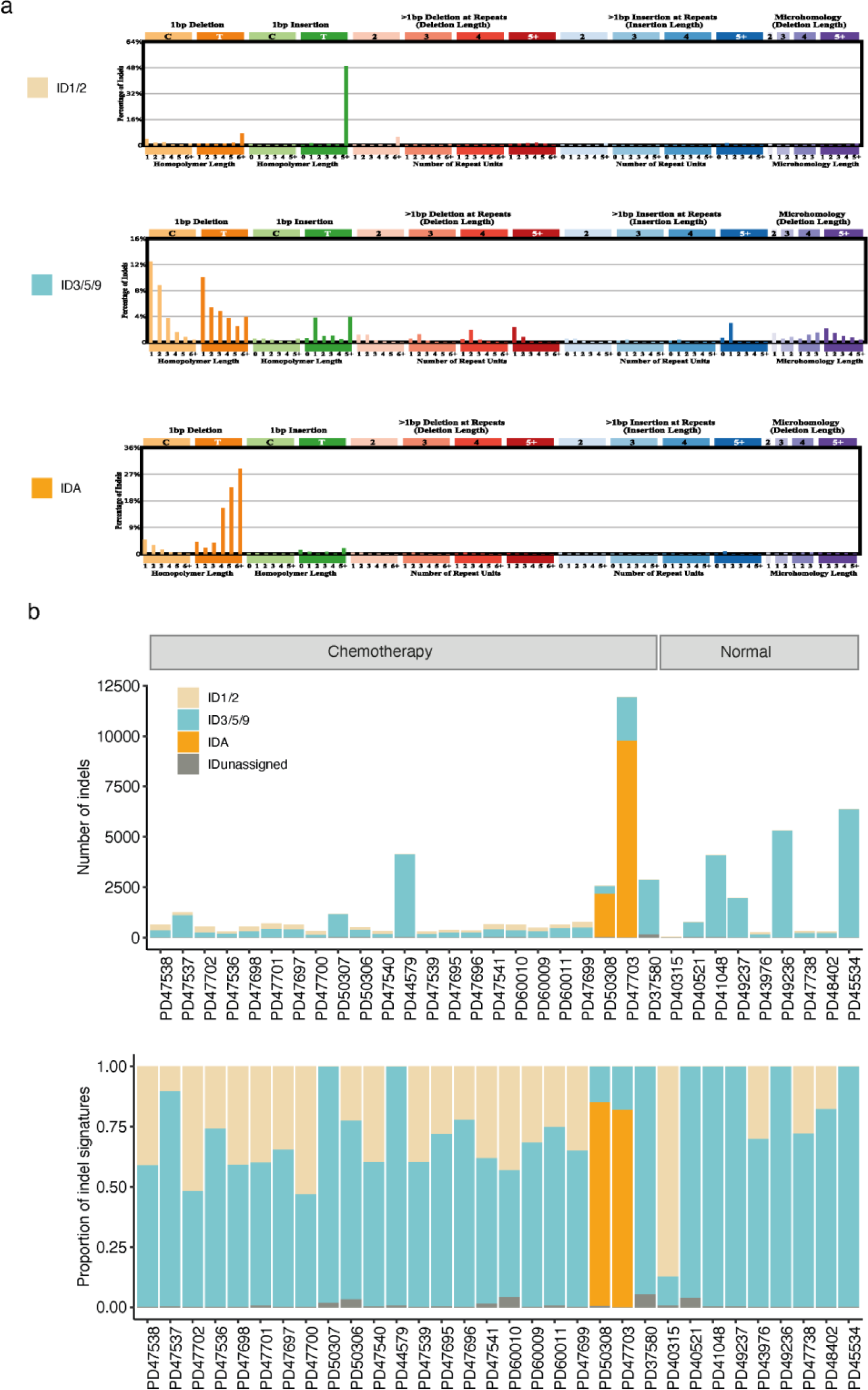
Indel signatures that are present in normal and chemotherapy exposed blood. a, Three indel signatures (ID1/2, ID3/5/9, IDA) were extracted by sigHDP. The context on the x-axis show the contributions of different types of indels, grouped by whether variants are deletions or insertions, the size of the event, the presence within repeat units and the sequence content of the indel. b, The proportion of indels and indels burden per mutational signatures across 22 chemotherapy exposed and 9 normal individuals, extracting using msigHDP (Methods). Each column represents samples from one individual. Signatures with the contribution <5% are considered as ‘unassigned’.

**Extended Data Fig. 4:**
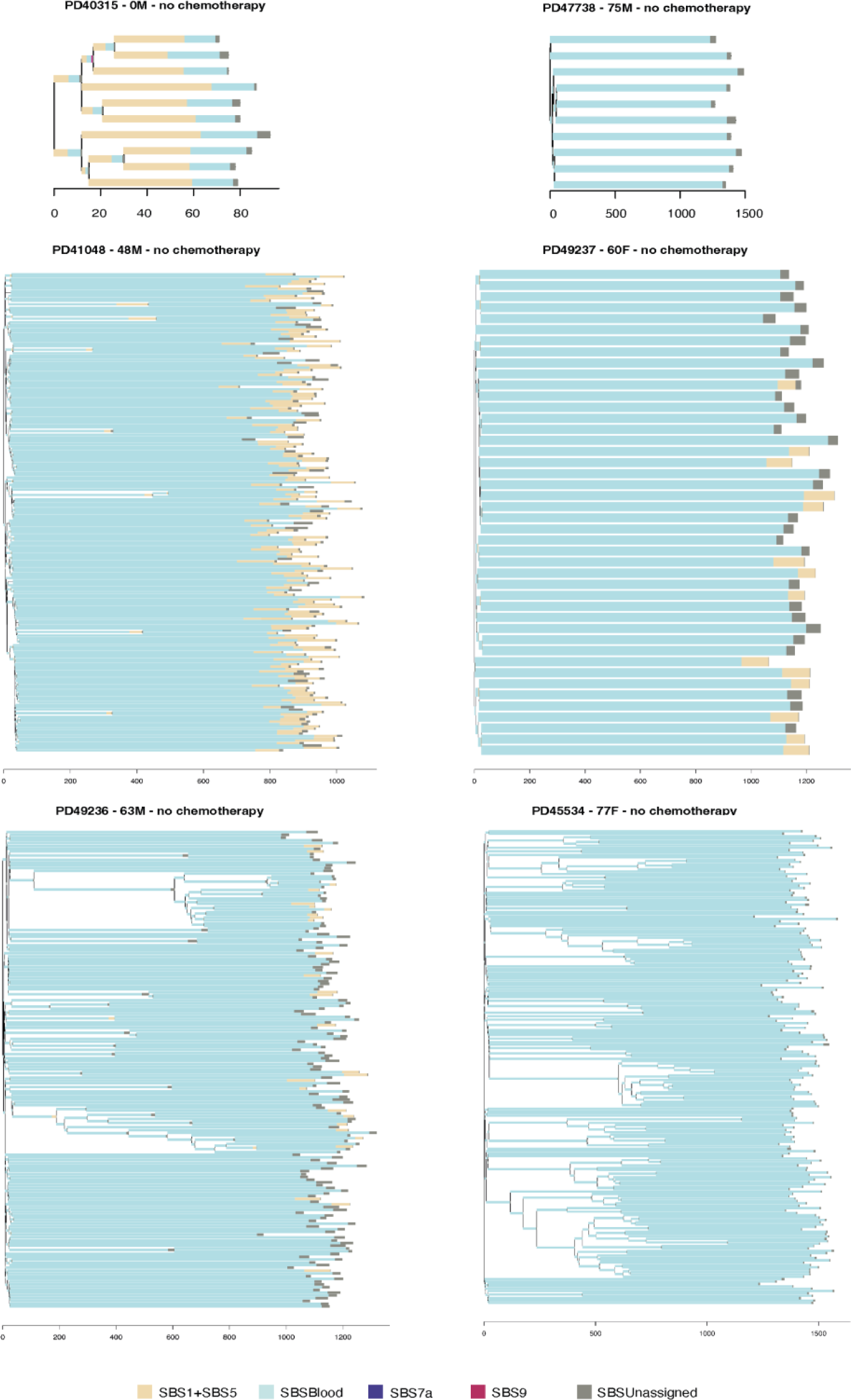
Phylogenetic trees and mutational signatures in normal individuals. Branch lengths correspond to SBS burdens. A stacked bar plot represents the SBS mutational signatures contributing to each branch with color code below the trees. SBSUnassigned indicates mutations that could not confidently be assigned to any reported signature.

**Extended Data Fig. 5:**
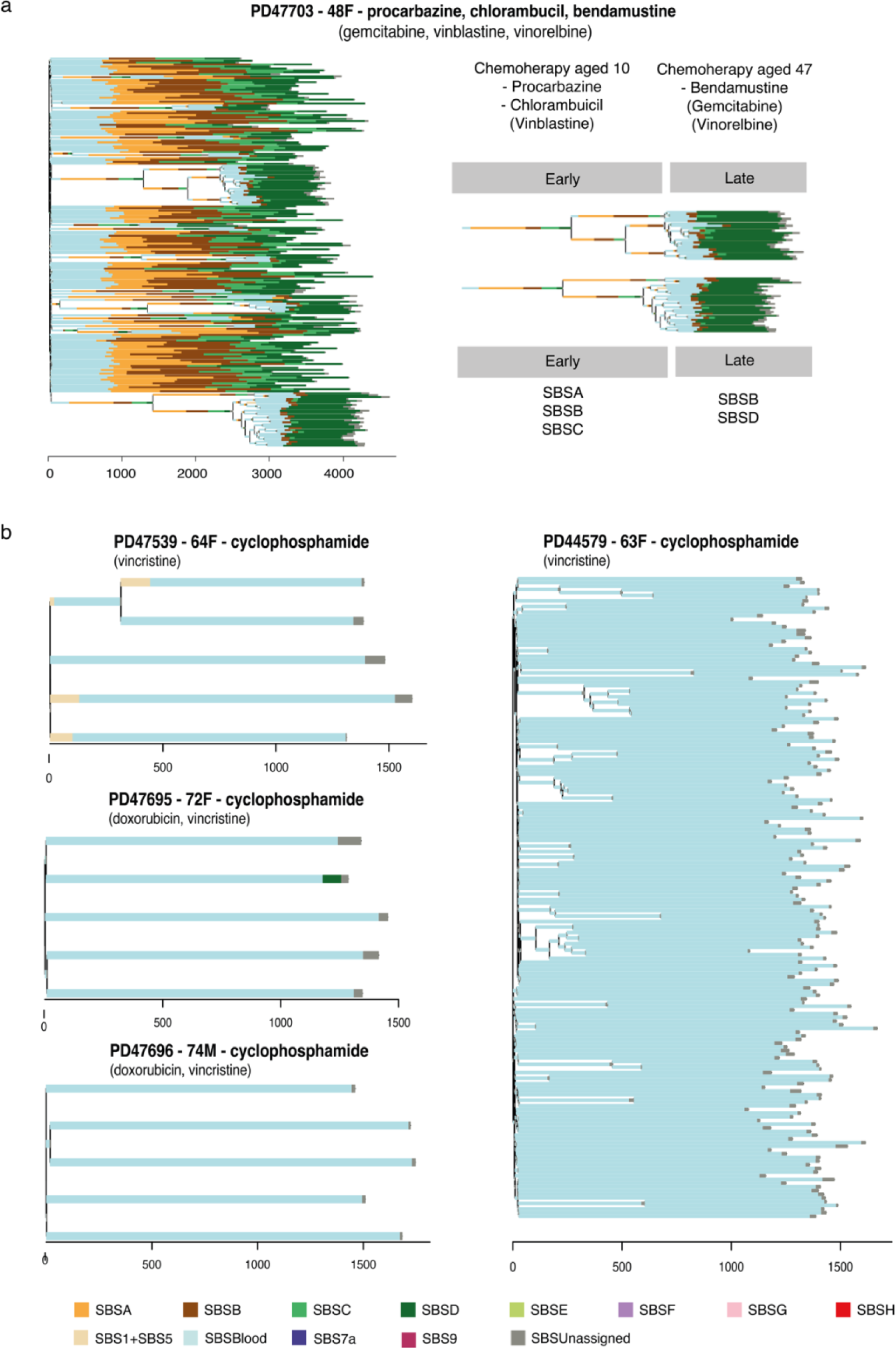
Phylogenetic trees and mutational signatures in individuals treated with alkylating agents. a, Phylogenetic tree of 48-year-old chemotherapy exposed female (PD47703). Branch lengths correspond to SBS burdens. A stacked bar plot represents the SBS mutationsal signatures contributing to each branch with color code below the trees. SBSUnassigned indicates mutations that could not confidently be assigned to any reported signature. She had been treated with chlorambucil and procarbazine at age 10 (early), and bendamustine at age 47 (late). b, Phylogenetic trees and SBS mutational signatures in individuals treated with cyclophosphamide.

**Extended Data Fig. 6:**
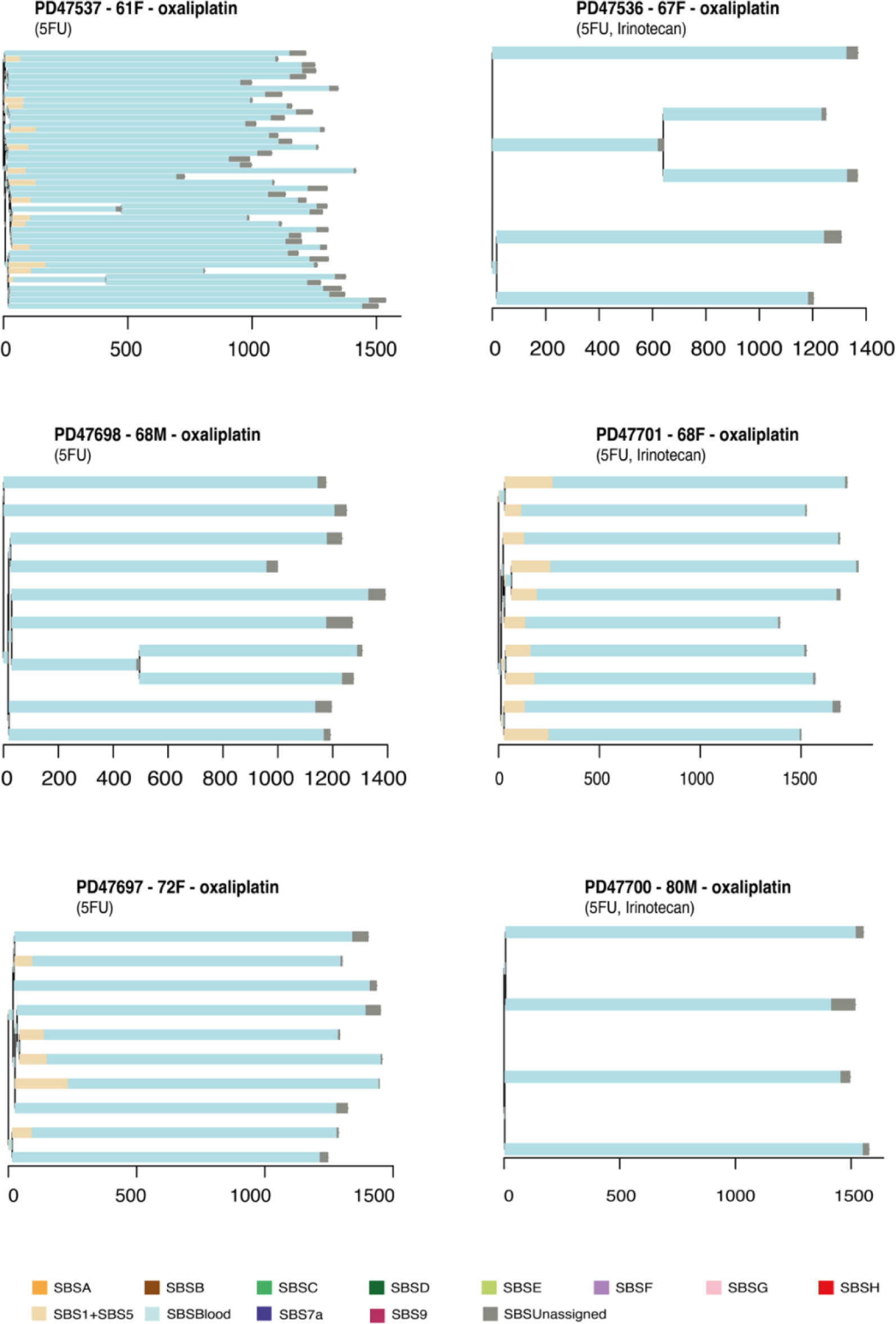
Phylogenetic trees and mutational signatures in individuals treated with oxaliplatin. Branch lengths correspond to SBS burdens. A stacked bar plot represents the SBS mutational signatures contributing to each branch with color code below the trees. SBSUnassigned indicates mutations that could not confidently be assigned to any reported signature.

**Extended Data Fig. 7:**
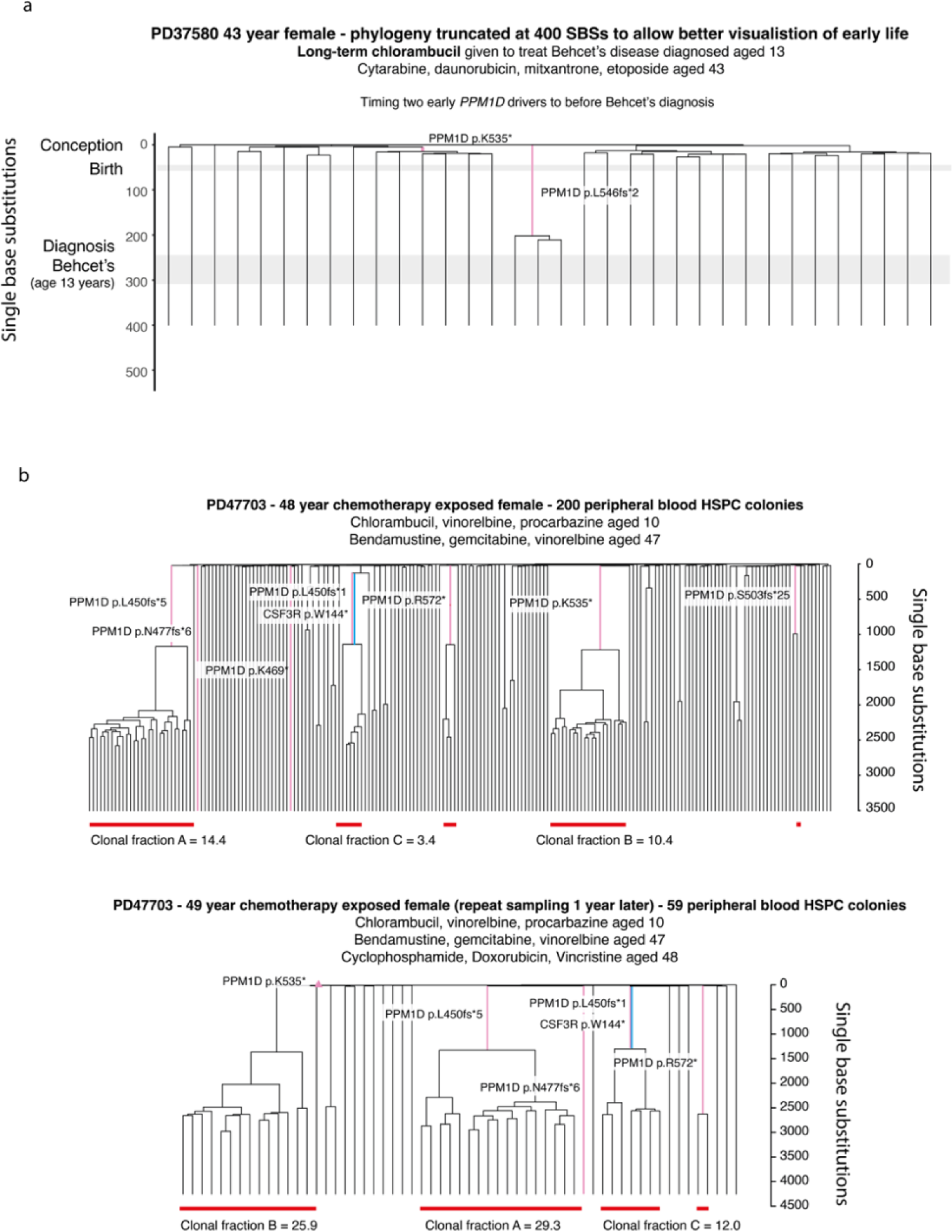
Annotated HSPC phylogenies for two chemotherapy treated individuals. Phylogenies were constructed for PD37580 (a) and PD47703 (b) individuals using shared mutation data and the algorithm MPBoot (Methods). In all phylogenies, branch lengths reflect the number of SBS mutations assigned to the branch. The y-axis represents the number of SBSs accumulating over time. Each tip on a phylogeny represents a single colony. Chemotherapy agents and the age of exposure to them are shown on top of the trees. a, PD37580 phylogeny of early life, truncated at 400 SBS mutations to allow better visualisation of the timing of acquisition of two early *PPM1D* mutations (pink). The number of mutations at age 13 was estimated using the linear mixed model described in Mitchell *et al* with 95% CI based on mutation burden being Poisson distributed as described in methods (241-306 single base subsitutions). b, Comparison of phylogenies created from peripheral blood samples taken from PD44703 one year apart. Pathogenic mutations in *PPM1D* have been highlighted (pink) to facilitate comparison of clone sizes at each timepoint. In addition a loss of function mutation in *CSF3R* has been highlighted (blue), which could also be contributing to loss of haematopoietic reserve and cytopenias. Red bars show the size of clonal fractions at each timepoint. Terminal branches have been adjusted for sequence coverage, and overall root-to-tip branch lengths have been normalized to the same total length (because all colonies were collected from a single time point).

**Extended Data Fig. 8:**
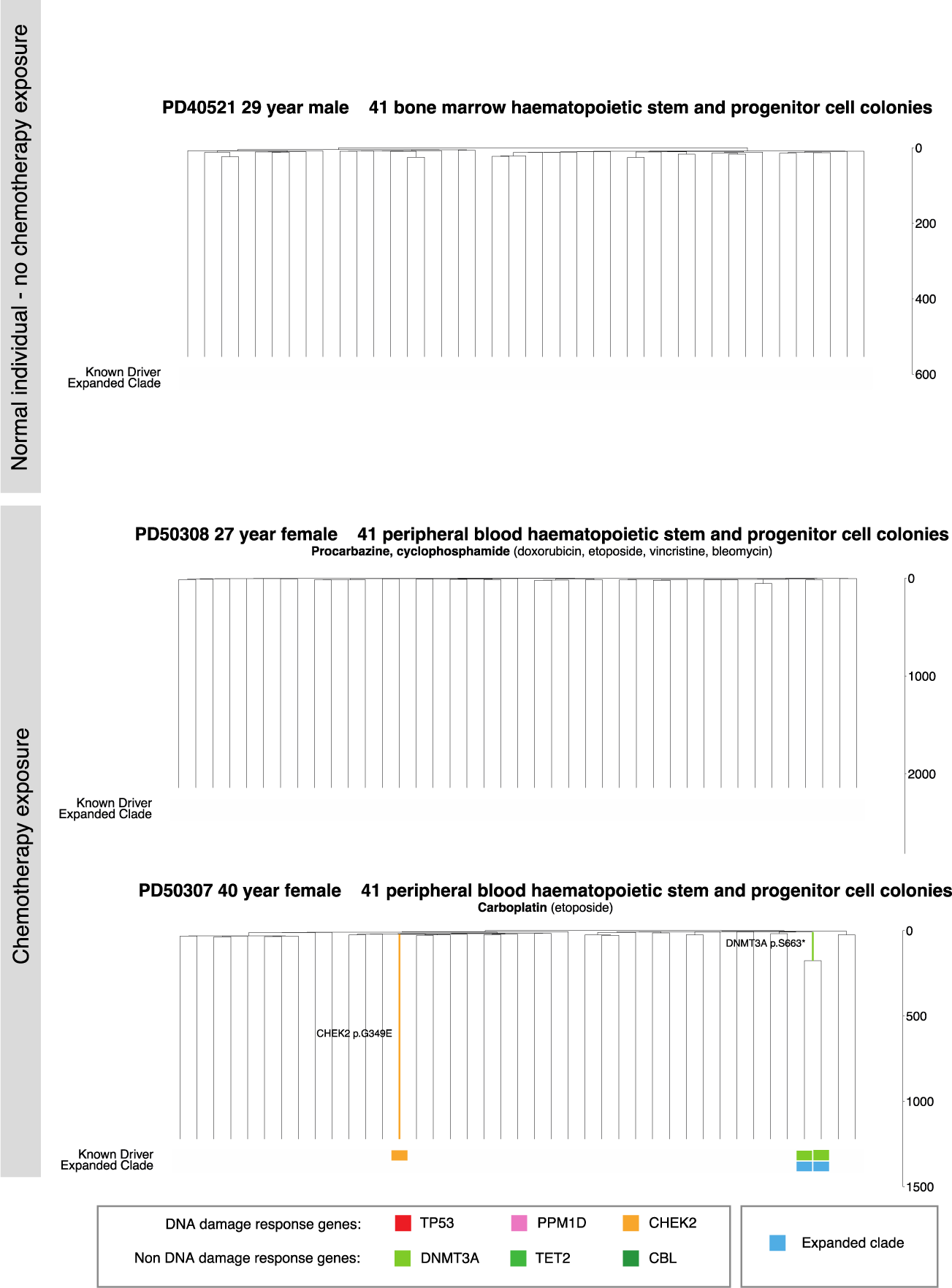
HSPC phylogenies for three young adult individuals. Phylogenies for one normal young adult individuals (top) and two young adult chemotherapy-treated individuals (bottom) were constructed using shared mutation data and the algorithm MPBoot (Methods). Branch lengths reflect the number of mutations assigned to the branch with terminal branches adjusted for sequence coverage, and overall root-to-tip branch lengths have been normalized to the same total length (because all colonies were collected from a single time point). The y-axis represents the number of SBSs accumulating over time. Each tip on a phylogeny represents a single colony, with the respective numbers of colonies of each cell and tissue type recorded at the top. Onto these trees, we have layered clone and colony-specific phenotypic information. We have highlighted branches on which we have identified known oncogenic drivers in one of 18 clonal haematopoiesis genes (Table S2) color-coded by gene. A heat map at the bottom of each phylogeny highlights colonies from known driver clades coloured by gene, and the expanded clades (defined as those with a clonal fraction above 1%) in blue.

**Extended Data Fig 9:**
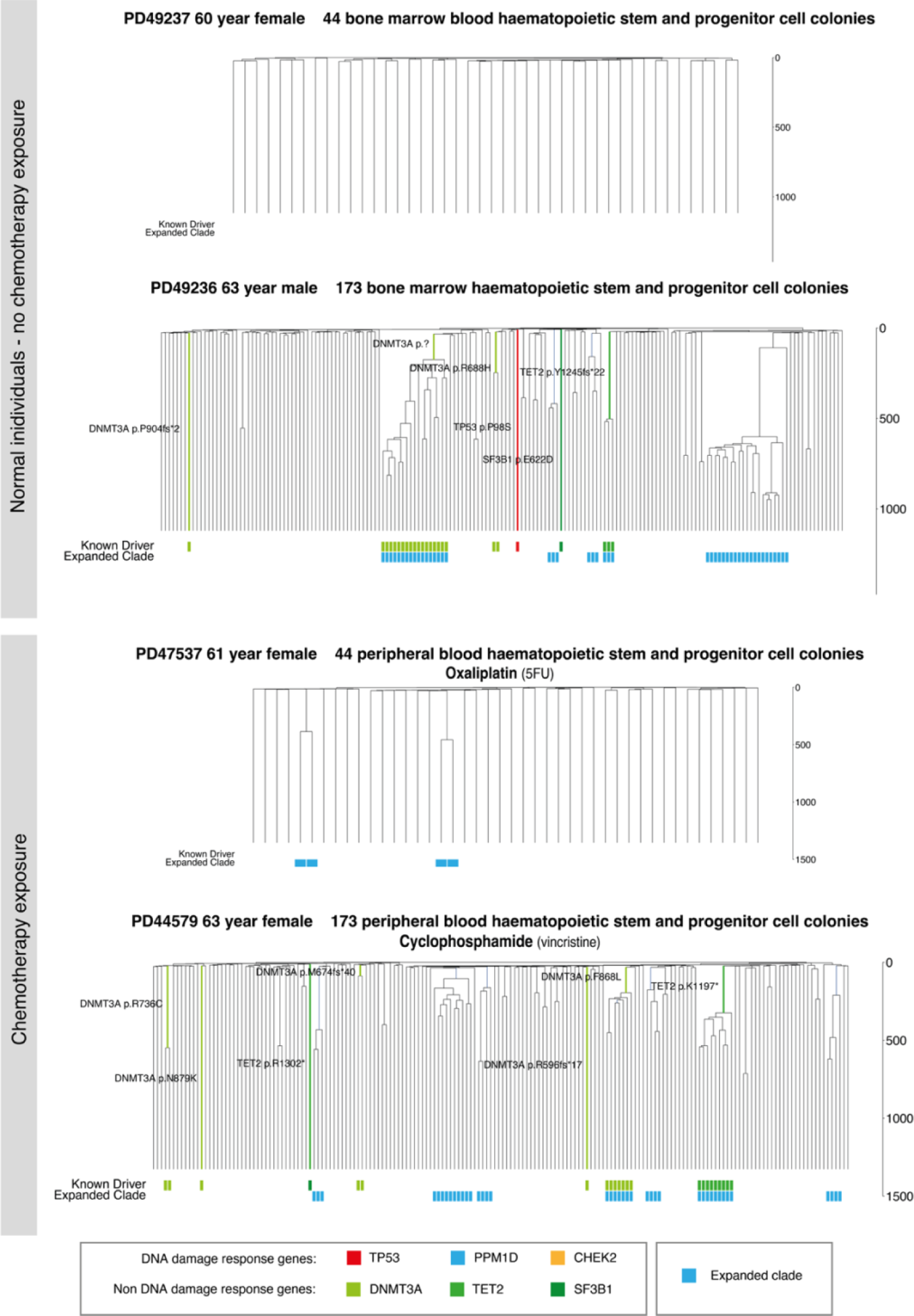
HSPC phylogenies for four older adult indiviudals. Phylogenies for two normal individuals (top) and two chemotherapy-treated individuals (bottom) were constructed using shared mutation data and the algorithm MPBoot (Methods). Branch lengths reflect the number of mutations assigned to the branch with terminal branches adjusted for sequence coverage, and overall root-to-tip branch lengths have been normalized to the same total length (because all colonies were collected from a single time point). The y-axis represents the number of SBSs accumulating over time. Each tip on a phylogeny represents a single colony, with the respective numbers of colonies of each cell and tissue type recorded at the top. Onto these trees, we have layered clone and colony-specific phenotypic information. We have highlighted branches on which we have identified known oncogenic drivers in one of 18 clonal haematopoiesis genes (Table S2) color-coded by gene. A heat map at the bottom of each phylogeny highlights colonies from known driver clades coloured by gene, and the expanded clades (defined as those with a clonal fraction above 1%) in blue.

## Supplementary information

## Supplementary Methods

### Data reporting

No statistical methods were used to predetermine sample size. The experiments were not randomized and the investigators were not blinded to allocation during experiments and outcome assessment.

### Samples

Blood or bone marrow samples from individuals un-exposed to chemotherapy were obtained from three sources: **1)** Stem Cell Technologies provided frozen mononuclear cells (MNCs) for the cord blood sample that had been collected with informed consent, including for whole genome sequencing (catalog #70007); all data previously published. **2)** Cambridge Blood and Stem Cell Biobank (CBSB) provided fresh peripheral blood samples taken with informed consent from two patients at Addenbrooke’s Hospital (NHS Cambridgeshire 4 Research Ethics Committee reference 07/MRE05/44 for samples collected pre-November 2019 and Cambridge East Ethics Committee reference 18/EE/0199 for samples collected from November 2019 onwards; all data previously published. **3)** Cambridge Biorepository for Translational Medicine (CBTM) provided frozen bone marrow +/- peripheral blood MNCs taken with informed consent from seven deceased organ donors. Samples were collected at the time of abdominal organ harvest (Cambridgeshire 4 Research Ethics Committee reference 15/EE/0152); data previously published from 4 individuals with new data generated from an additional 2 individuals (PD49236 and PD49327).

Blood samples from individuals previously exposed to chemotherapy were obtained from two sources: **1)** Cambridge Blood and Stem Cell Biobank (CBSB) provided fresh peripheral blood samples taken with informed consent from 22 patients at Addenbrooke’s Hospital (NHS Cambridgeshire 4 Research Ethics Committee reference 07/MRE05/44 for samples collected pre-November 2019 and Cambridge East Ethics Committee reference 18/EE/0199 for samples collected from November 2019 onwards; all unpublished data. One chemotherapy exposed individual, PD44703, had two samples taken at timepoints a year apart. All others were sampled at a single timepoint. **2)** Baylor College of Medicine provided single cell-derived haematopoietic colonies from bone marrow taken following informed consent from 1 patient from MD Anderson Cancer Centre; Research Ethics Committee of the University of Texas MD Anderson Cancer Centre Institutional Review Board reference PA12-0305 (genomic analysis protocol) and LAB01-473 (laboratory protocol).

Details of the individuals studied and the samples they provided are listed in **Fig. 1a**, with additional information in **Table S1**.

### Isolation of MNCs from fresh peripheral blood samples

Whole blood was diluted 1:1 with PBS, after which mononuclear cells (MNCs) were isolated using lymphoprep^TM^ density gradient centrifugation (STEMCELL Technologies. Red cell lysis was performed on the MNC fraction using 1 incubation at 4°C for 15 mins with RBC lysis buffer (BioLegend).

### Single-cell colony expansion *in vitro* – liquid culture (unexposed samples)

For all the unexposed samples normal samples and the PD44703 second timepoint sample, single-cell colony expansion *in vitro* was undertaken in liquid culture, exactly as previously described^1^.

Peripheral blood and cord blood MNC samples underwent CD34+ selection using the EasySep human whole blood CD34 positive selection kit (STEMCELL Technologies), with only a single round of magnetic selection. Bone marrow MNCs were not CD34 selected prior to cell sorting.

MNC or CD34 enriched samples were stained (30 minutes at 4°C) in PBS/3%FBS containing the following antibodies: CD3/FITC, CD90/PE, CD49f/PECy5, CD38/PECy7, CD33/APC, CD19/A700, CD34/APCCy7, CD45RA/BV421 and Zombie/Aqua (**Table S3**). Cells were then washed and resuspended in PBS/3%FBS for cell sorting. Either a BD Aria III or BD Aria Fusion cell sorter (BD Biosciences) was used to sort ‘HSC/MPP’ pool cells (Lin-, CD34+, CD38-, CD45RA-) at the NIHR Cambridge BRC Cell Phenotyping hub. **Supplementary Fig. 1** illustrates the gating strategy used.

**Supplementary Fig. 1:**
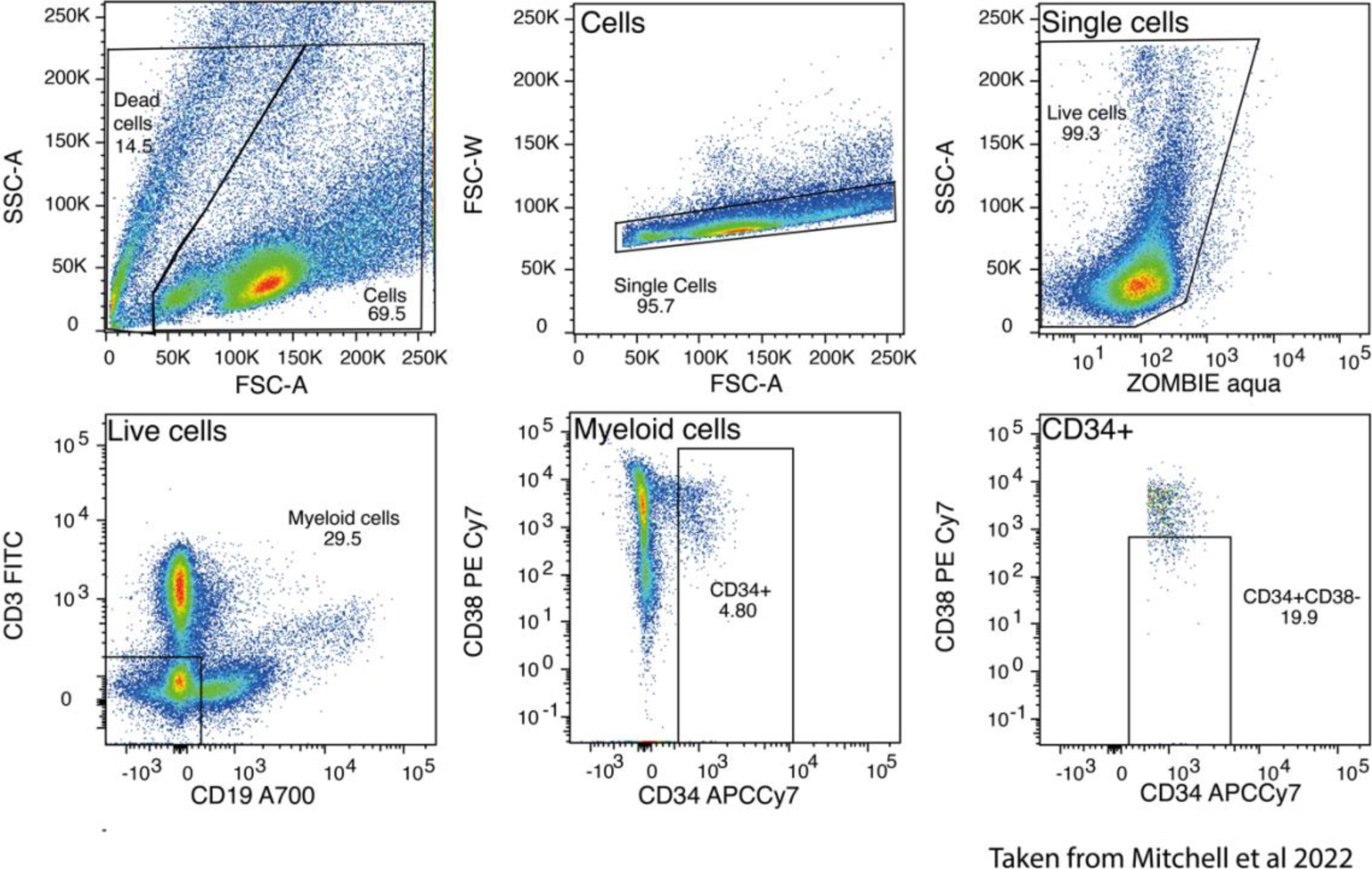
Flow-sorting strategy for single HSPCs (CD34+CD38-) cells. **A**, Sorting of single human HSPCs from cord blood, peripheral blood and bone marrow. Cells were stained with the panel of antibodies in **Table S3** then single HSPCs were index sorted according to the strategy depicted into individual wells of 96 well plates.

Single phenotypic ‘HSPC’ cells were index sorted into single wells of 96 well plates containing StemPro media (Stem Cell Technogies), StemPro Nutrients (0.035%, Stem Cell Technologies), L-Glutamine (1%, ThermoFisher), Penicillin-Streptomycin (1%, ThermoFisher) and cytokines (SCF, 100 ng/ml; FLT3, 20 ng/ml; TPO, 100 ng/ml; EPO 3 ng/ml; IL-6, 50 ng/ml; IL-3, 10 ng/ml; IL-11, 50 ng/ml; GM-CSF, 20 ng/ml; IL-2 10 ng/ml; IL-7 20 ng/ml; lipids 50 ng/ml), and expanded into colonies. Cells were incubated at 37°C and the colonies that formed were topped up with 50μl StemPro media plus supplements at 14 +/- 2 days as necessary. At 21 +/- 2 days colonies were harvested. DNA extraction was performed using either the DNEasy 96 blood and tissue plate kit (Qiagen) or the Arcturus Picopure DNA Extraction kit (ThermoFisher) as per the manufacturer’s instructions.

### Single-cell colony expansion *in vitro* – Methocult (chemotherapy-exposed samples)

For the chemotherapy-exposed peripheral blood samples, single-cell colonies were expanded in Methocult (H4435 or H4034, STEMCELL Technologies). MNCs were plated at a density of 7.5-45-x10^4^/ml in Methocult and incubated at 37C for 14 days. The cell suspensions were made up in StemSpan II (STEMCELL technologies) before being mixed thoroughly with MethoCult and plated into a SmartDish (STEMCELL technologies). Individual BFU-E or CFU-GM colonies were picked into 17ul of proteinase K (PicoPure DNA extraction kit, Fisher Scientific-each vial lyophilised proteinase k resuspended in 130ul reconstitution buffer) and incubated 65C for 6hrs, 75C for 30mins to extract DNA in preparation for sequencing.

Previous studies have shown that there is no difference in mutation burden between HSCs and haematopoietic progenitor cells^1^ and that there is only a mutation burden difference of ∼30 SBS mutations between HSCs and mature granulocytes^2^.

### Whole genome sequencing of colonies

Whole genome sequencing libraries were prepared from 1-5ng of extracted DNA from each colony using a low input enzymatic fragmentation-based library preparation method^3,4^. Whole genome sequencing was performed on the NovaSeq platform (Illumina). 150bp paired end reads were aligned to the human reference genome (NCBI build37) using BWA *mem*.

### Single-base-substitution and indel calling

The method for substitution calling involves three main steps: mutation discovery, filtering, and genotyping, as describe in previous paper (ref-Lee-Six, H. et al., https://doi.org/10.1038/s41586-019-1672-7).

Mutation discovery is initiated using the CaVEMan algorithm^5^, configured with copy-number settings of major copy number 5 and minor copy number 2 for normal clones to maximize sensitivity. An unmatched normal sample is utilized to prevent the misclassification of embryonic mutations as germline mutations, crucial for accurate phylogenetic analysis.

Various filters are applied to the data. Filtering against a panel of 75 unmatched normal samples helps eliminate common single-nucleotide polymorphisms (SNPs). Additional filters target mapping artefacts associated with BWA-MEM alignment, setting thresholds such as requiring a median alignment score ≥140 and less than half of reads to be clipped. Fragment-based statistics are employed to prevent calling variants supported by a low number of fragments. Variants are further annotated and filtered based on fragment coverage, number of fragments supporting the variant, fragment-based allele fraction.

The genotyping stage involves creating a pile-up of all samples from an individual, counting mutant and wild-type reads. Stringent criteria, including a variant allele frequency (VAF) >0.2, depth >7, and ≥4 mutant reads, are employed for mutation calling. Positions with insufficient data or conflicting information across samples are marked as not applicable (NA) for tree construction. Germline positions are identified based on consistent presence or NA status across samples from an individual.

### Insertion/ deletion calling

Indels were called with the Pindel argorithm^6^ using a matched normal. The same dataset-specific filters used for substitutions as described above were also applied to indels. Subsequently, indels were genotyped, requiring a VAF >0.2, a minimum depth of 10, and support from at least 5 mutant reads.

### Structural variant and copy-number calling

Structural variants (SVs) were detected using GRIDSS8, confirmed visually and by adhering to the expected phylogenetic distribution based on single-base substitutions (SBS). SVs larger than 1kb with QUAL >=250 and those smaller than 30kb with QUAL >=300 were retained. Additionally, SVs required support from at least four discordant and two split reads, with a standard deviation of alignment positions > five being filtered out. A panel of normal samples (n=350) was incorporated to the GRIDSS panel to eliminate potential germline SVs and artefacts.

Autosomal copy number aberrations (CNAs) were identified using ASCAT (Allele-Specific Copy number Analysis of Tumours)^5^. The matched normal sample with coverage > 15X and no Y loss was selected for as for call structure variants. The in-house algorithm BRASS (Breakpoint AnalySiS)^6^ was used to call CNAs on sex chromosomes by generating read count information across 500bp segments. Y loss was determined by comparing X and Y chromosome coverage means, validated through visual inspection of read depth.

## Additional variant filtering steps

### Larger dataset containing samples sequenced at lower sequencing depth

For creation of the larger phylogenies (subset of 6 chemotherapy and 5 unexposed individuals with more than 40 sequenced colonies), a binomial filtering strategy could be applied as previously described^1^. (https://github.com/emily-mitchell/normal_haematopoiesis/2_variant_filtering_tree_building/scripts/).

### Small dataset containing samples sequenced at relatively high sequencing depth

For the subset of colonies sequenced at highest depth (4-10 colonies per individual), we were unable to use binomial filtering approaches due to low sample number. Instead the following filters were applied to the data using a custom R script: 1) Variants present in more than half the samples from an individual were removed as being most likely germline. 2) Variants were only called as present in a given sample if they had 2 or more supporting reads and were present at a variant allele fraction ≥ 0.2 in autosomes or ≥ 0.4 for sex chromosomes. 3) High and low depth sites with a mean depth more than 50 or less than 8 across all samples from an individual were removed.

This variant filtering approach was validated using samples from the normal individuals, in whom both the binomial and non-binomial filtering strategies were applied to the same samples, giving comparable results (**Supplementary fig. 2**). This dataset, comparable across all the individuals in the study, was used for analysis of SBS and indel mutation burdens.

**Supplementary figure 2:**
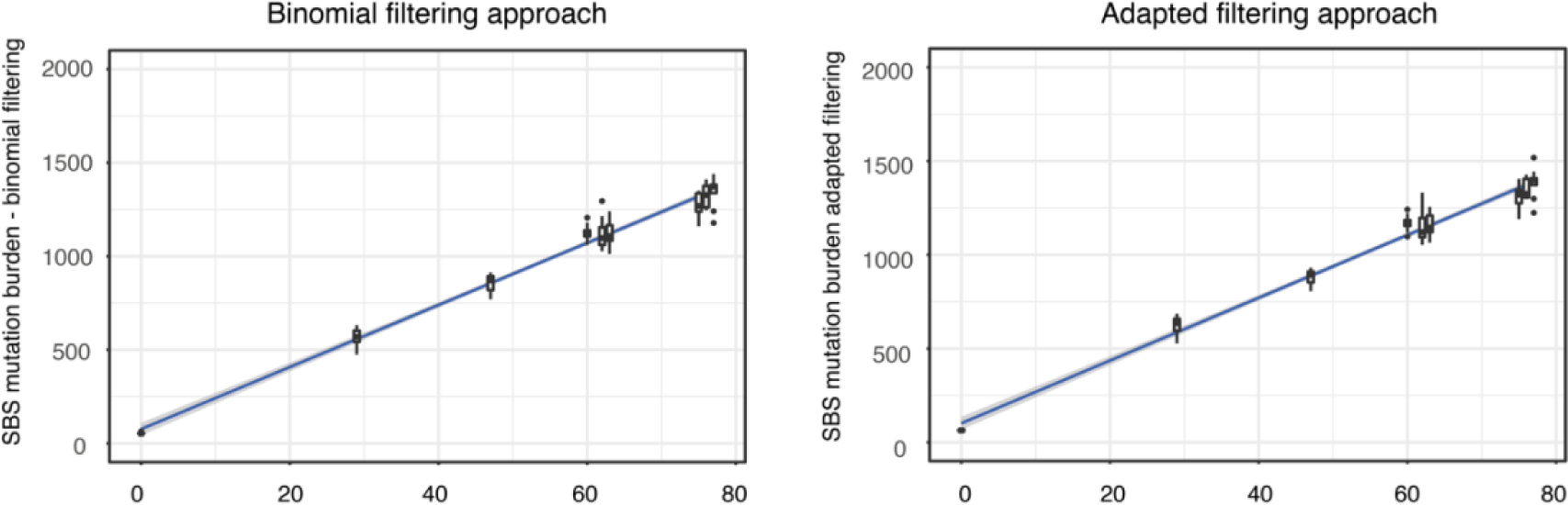
Validation of variant filtering approach. **a**, SNV mutation burden of 10 samples per normal individual with variants filtered using the binomial filtering strategy that made use of several hundred other samples sequenced from the individual (left). SNV mutation burden of the same 10 samples per normal individual with variants filtered using the adapted filtering strategy that required only 10 samples per individual (right).

### Filtering at the colony level

We removed a total of 96 colonies from the dataset of 931 previously unpublished colonies: 32 for being technical duplicates, 29 for showing evidence of non-clonality or contamination and 23 due to low coverage. Visual inspection of the VAF distribution plots and a peak VAF threshold of < 0.4 was used (after the removal of *in vitro* variants) to identify colonies with evidence of non-clonality.

### Mutation burden analysis

Due to the difficulty in correcting for sequencing depth when only a small number of samples is sequenced per individual, SBS and indel burden analysis was performed on raw data from the subset of chemotherapy and unexposed samples sequenced at relatively high depth (4-10 samples per individual; mean coverage 23X, range 13X-33X). We have previously shows that sequencing depth has little impact on SBS mutation burden over this higher range^1^. There were minor differences in sequencing depth when comparing the chemotherapy and normal cohorts or comparing sequencing depth by chemotherapy exposure that would not be expected to impact the interpretation of the results presented (**Extended Fig. 1**).

### Construction of phylogenetic trees

*MPBoot*, a maximum parsimony tree approximation method^7^, was used to build and annotate phylogenetic trees of the relationships between their sampled HSPCs as previously described^1^.

The key steps to generate the phylogenies shown in **Figures 5 and Extended Figs. 8 and 9** are as follows:

1. <u>Generate a ‘genotype matrix’ of mutation calls for every colony within a donor</u> – Our protocol, based on whole genome sequencing of single-cell-derived colonies, generates consistent and even coverage across the genome, leading to very few missing values within this matrix (ranging from 0.005 – 0.034 of mutated sites in a given colony across different donors within our cohort). This generates a high degree of accuracy in the constructed trees.
2. <u>Reconstruct phylogenetic trees from the genotype matrix</u> – This is a standard and well-studied problem in phylogenetics. The low fraction of the genome that is mutated in a given colony (<1/million bases) coupled with the highly complete genotype matrix mean that different phylogenetics methods produce reassuringly concordant trees. We used the *MPBoot* algorithm for the tree reconstruction, as it proved both accurate and computationally efficient for our dataset.
3. <u>Correct terminal branch lengths for sensitivity to detect mutations in each colony</u> *–* The trees generated in the previous step have branch lengths proportional to the number of mutations assigned to each branch. For the terminal branches, which contain mutations unique to that colony, variable sequencing depth can underestimate the true numbers of unique mutations, so we correct these branch lengths for the estimated sensitivity to detect mutations based on genome coverage.
4. <u>Make phylogenetic trees ultrametric</u> – After step 3, there is little more than Poisson variation in corrected mutation burden among colonies from a given donor. Since these colonies all derived from the same timepoint, we can normalise the branch lengths to have the same overall distance from root to tip (known as an ultrametric tree). We used an ‘iteratively reweighted means’ algorithm for this purpose.
5. <u>Scale trees to chronological age</u> – Since mutation rate is constant across the human lifespan, we can use it as a ‘molecular clock’ to linearly scale the ultrametric tree to chronological age.
6. <u>Overlay phenotypic and genotypic information on the tree</u> – The tip of each branch in the resulting phylogenetic tree represents a specific colony in the dataset, meaning that we can depict phenotypic information about each colony underneath its terminal branch (the coloured stripes along the bottom of **Fig. 5 and Extended Figs. 8 and 9**). Furthermore, every mutation in the dataset is confidently assigned to a specific branch in the phylogenetic tree. This means that we can highlight branches on which specific genetic events occurred (such as *DNMT3A* or other driver mutations).

More detailed information on these steps is provided below:

*MPBoot*, a maximum parsimony tree approximation method^7^, was used to build phylogenetic trees of the relationships between the sampled cells. Variants were genotyped as ‘present’ (coded as 1) in a sample if 2 or more variant reads supported the variant. Variants were genotyped as ‘absent’ (coded as 0) in a sample if 0 variant reads were present at a given site and depth at that site was 6 or more. Sites that did not fall into either of the above categories were marked as ‘unknown’ (coded as 0.5). In all cases only a small minority of sites (< 5%) were categorised as ‘unknown’ or ‘missing data’ as shown in the table below.

The genotype matrix of shared variants was converted to a ‘DNA string’ for each sample with ‘W’ representing a ‘wildtype’ position, ‘V’ a ‘variant’ position and ‘?’ representing ‘unknown’. The DNA strings were then used as the input for *MPBoot*, which outputs unscaled trees with uninformative branch lengths. We explicitly added a ‘dummy sample’ (called “Ancestral”) into the DNA strings that *MPBoot* used, which has non-mutant genotypes across all sites i.e. representing the genotypes of the reference genome. After tree construction the ‘ancestral’ branch was dropped prior to downstream analyses.”A maximum likelihood approach and the original count data was then used to assign each mutation in an individual’s dataset to a branch in their *MPBoot* generated phylogenetic tree (https://github.com/NickWilliamsSanger/treemut). Tree edge lengths were then made proportional to the number of mutations assigned to the branch.

The sensitivity of mutation calling in each sample was used to correct phylogeny branch lengths for sequencing coverage. Sensitivity was calculated as the fraction of known germline variants identified by CaVEMan in a specific sample. Mutation burden was corrected by multiplying the number of variants by 1/sensitivity for private branches. The sensitivity was adjusted to allow for the higher sensitivity on shared branches due to multiple samples containing the variant. Specifically, sensitivity was assessed by measuring the ability of the mutation-calling algorithms to detect heterozygous germline single nucleotide polymorphisms (SNPs) in each sample. Heterozygous SNPs should have the same VAF distribution and sensitivity as true somatic mutations. For private branches, the SNV component of branch lengths was scaled according to:

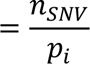

Where *n_cSNV_* is the corrected number of SNVs in sample *i*, *n_SNV_* is the uncorrected number of SNVs called in sample *i* and *p_i_* is the proportion of germline SNPs called by the Caveman algorithm in sample *i*.

For shared branches, it was assumed that (1) the regions of low sensitivity were independent between samples, (2) if a somatic mutation was called in at least one sample within the clade, it would also be correctly called (or ‘rescued’) in other samples in the clade (even in lower sensitivity samples). Shared branches were therefore scaled according to:

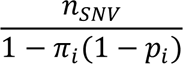

Where the product is taken for 1 − *p_i_* for each sample *i* within the clade. However, both of these assumptions will not hold true in all cases. Firstly, regions with low coverage are not randomly distributed, with some genomic regions likely to have low coverage in multiple samples. Secondly, while many mutations will be ‘rescued’ in subsequent samples once they have been called in a first sample - because the *treemut* algorithm for mutation assignment uses original read count data, meaning that even a single variant read in a subsequent sample is likely to result in the mutation being correctly assigned - this will not be true in every case. Some samples with very low coverage have 0 variant reads at a given site will by chance. In this situation, a mutation may not be correctly placed. While these factors may lead to an under-correction of shared branches, this approach provides a reasonable approximation. Corrected SNV burdens for each sample can then be calculated as the sum of corrected ancestral branch lengths back to the root of the phylogeny.

The phylogenies were then made ultrametric (or linearised) using a previously published bespoke algorithm to make all branch lengths equal^1^. Starting from the root of the tree and moving progressively towards each tip, the fraction of time for the given shared branch is calculated as the fraction of remaining time times the number of mutations on the given shared branch divided by the mean number of mutations of all descendants from that shared branch. The function is called recursively, updating the fraction of remaining time, as the algorithm moves from root to tip. This algorithm therefore has the property that the most confident timings (nodes near the root) are defined first, anchoring the timings of subsequent, less confident nodes.

Additional information in the form of driver mutations was then overlaid on the final ultrametric version of to generate the final phylogenies depicted in **Fig. 5 and Extended Figs. 8 and 9**.

To estimate the number of somatic mutations that may have already been acquired by PD37580 by age 13 (prior to commencing chlorambucil), we used the linear mixed model defined in Mitchell et al^1^. This model estimates an intercept of 54.57 (ie the mean number of somatic mutations present at birth), with a slope of 16.832 representing the mean number of somatic mutations acquired each year of life. This results in an expected mean somatic mutation burden of 273 at age 13. Assuming this mutation burden is Poisson distributed provides a 95% prediction interval of 241-306.

### Analysis of driver variants

Variants identified were annotated with VAGrENT (Variation Annotation GENeraTor) (https://github.com/cancerit/VAGrENT) to identify protein coding mutations and putative driver mutations in each dataset. **Table S4** lists the 18 genes we have used as our top clonal haematopoiesis genes (those identified by Fabre *et al* as being under positive selection in a targeted sequencing dataset of 385 older individuals, with CHEK2 added as being an additional gene commonly under positive selection in chemotherapy exposed individuals). ‘Oncogenic’ mutations (as assessed by EM) are shown in **Figure 5 and Extended Figs. 8 and 9**.

## Bulk cell sorts for Nanoseq sequencing (chemotherapy-exposed samples)

### Chemotherapy-exposed samples

Mononuclear cells were stained for 30 minutes at 4C in PBS/3%FCS containing the following antibodies: Zombie Aqua, CD3 APC, CD19 AF700, CD45RA PerCPCy5.5, CCR7 BV711, CD14 BV605. Cells were then washed and resuspended in PBS/3%FBS for cell sorting. Either a BD Aria III or BD Aria Fusion cell sorter (BD Biosciences) was used to sort various mature cell compartments (B cells, T naive cells, T memory cells, and monocytes) at the NIHR Cambridge BRC Cell Phenotyping hub. For each cell type ∼40,000 cells were sorted into Eppendorf tubes containing 50 μl PBS. Further details in **Table S4** and **Supplementary Fig. 3**.

**Supplementary figure 3:**
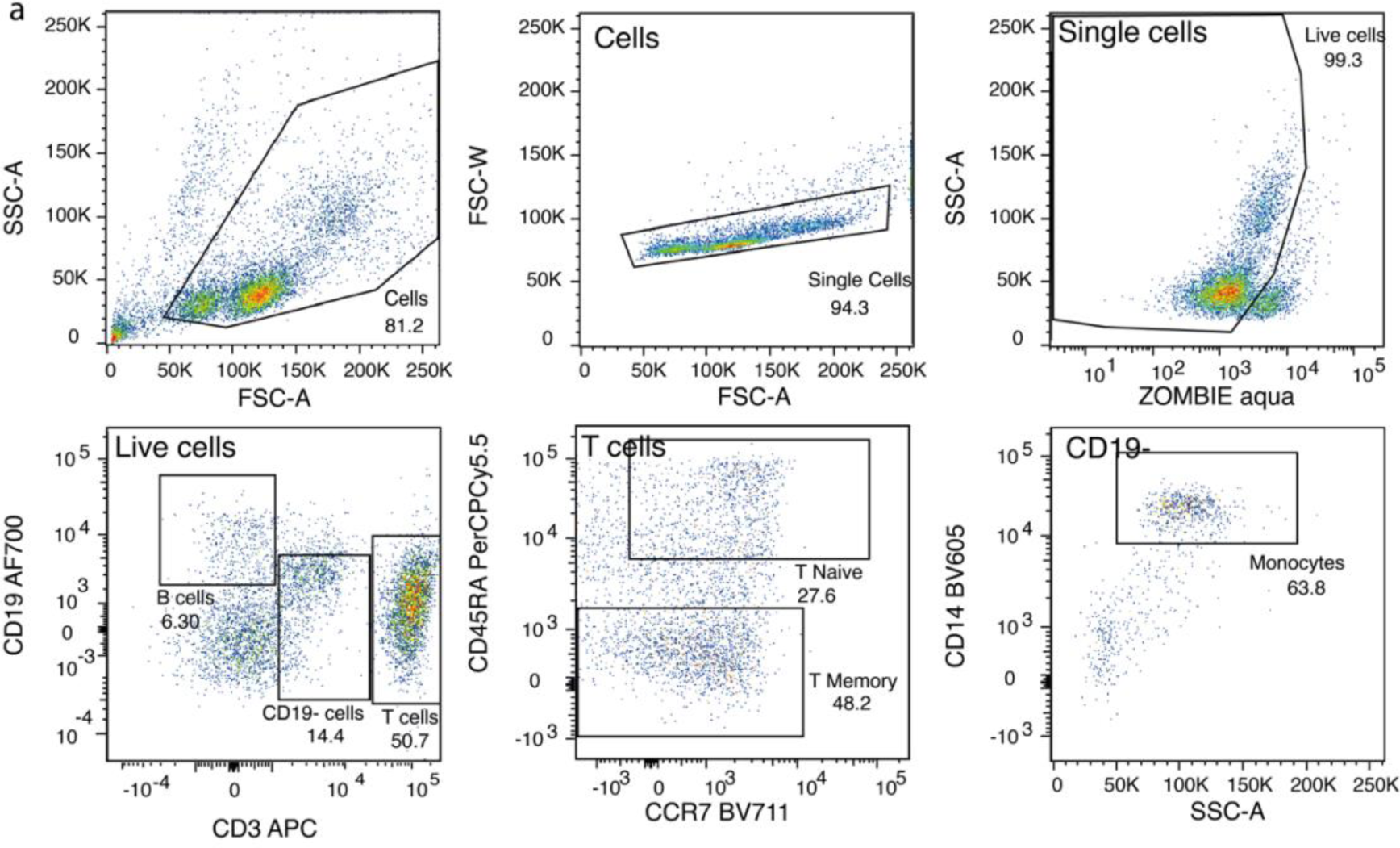
Flow-sorting strategy for single mature blood cells from chemotherapy exposed individuals. **a**, Sorting of single human mature blood cells from peripheral blood. Cells were stained with the panel of antibodies in **Table S4** then single cells were bulk sorted according to the strategy depicted into individual Eppendorf tubes.

### Unexposed normal samples

Mononuclear cells were stained for 30 minutes at 4C in PBS/3%FCS containing the following antibodies: CD3 APC, CD4 BV785, CD8 BV785, CD14 BV605, CD19 AF700, CD20 PE Dazzle, CD27 BV421, CD34 APC-Cy7, CD38 FITC, CD45RA PerCPCy5.5, CD56 PE, CCR7 BV711, IgD PECy7, Zombie Aqua. Cells were then washed and resuspended in PBS/3%FBS for cell sorting. Either a BD Aria III or BD Aria Fusion cell sorter (BD Biosciences) was used to sort various mature cell compartments (B cells, CD4+T naive cells, CD4+T memory cells, CD8+ T naïve cells, CD8+ T memory cells and monocytes) at the NIHR Cambridge BRC Cell Phenotyping hub. For each cell type ∼40,000 cells were sorted into Eppendorf tubes containing 50 μl PBS. Further details in **Table S5** and **Supplementary Fig. 4**:

**Supplementary figure 4:**
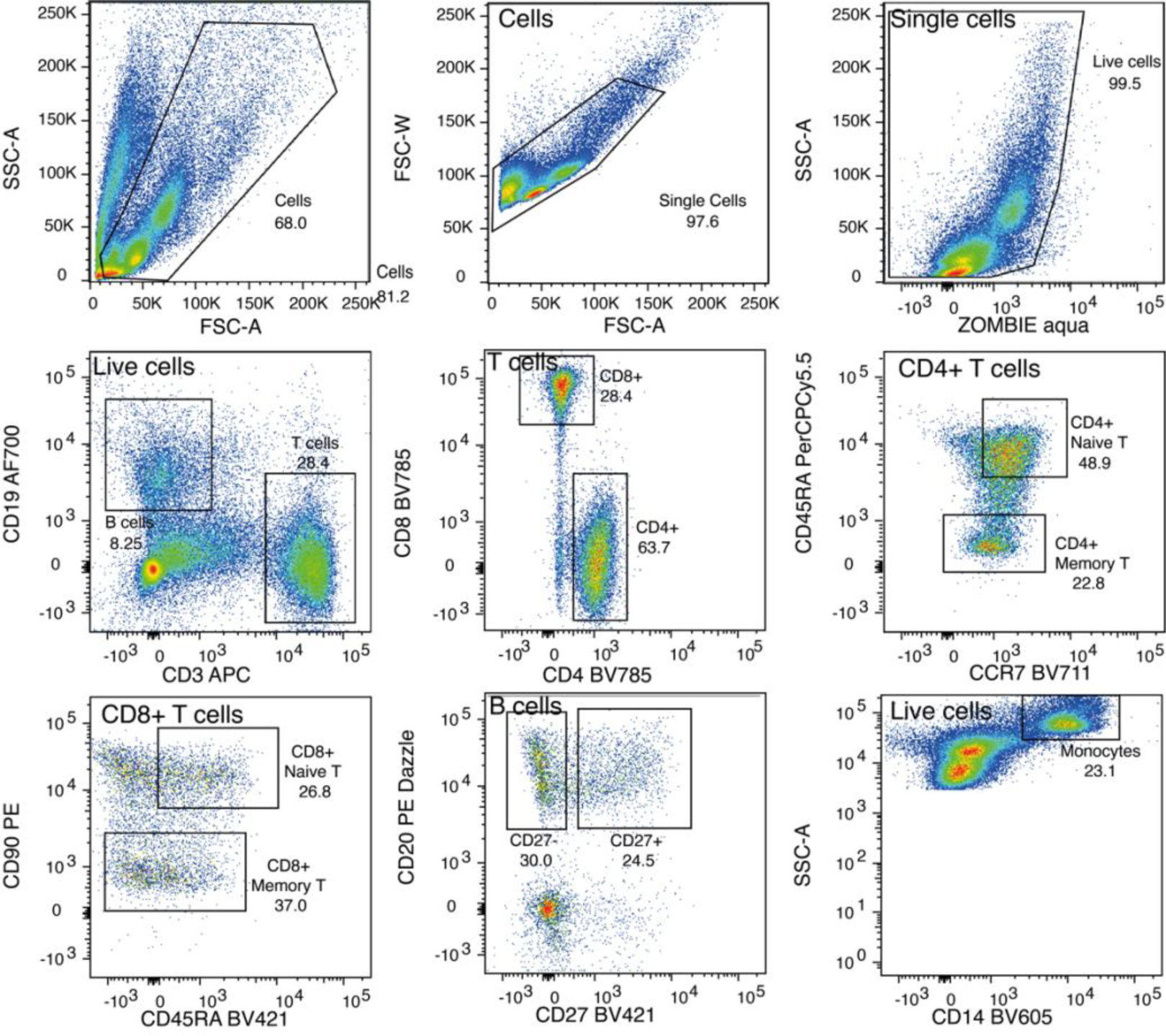
Flow-sorting strategy for single mature blood cells from unexposed normal individuals. **a**, Sorting of single human mature blood cells from peripheral blood. Cells were stained with the panel of antibodies in **Table S5** then single cells were bulk sorted according to the strategy depicted into individual Eppendorf tubes.

### DNA extraction from bulk mature cell sorts

Approximately 40,000 cells of each mature cell type from the above sorts sort (suspended in 200ul PBS) were added to single wells of a 96 well PCR plate and centrifuged. Pellets were resuspended in 17ul proteinase K (PicoPure DNA extraction kit, Fisher Scientific; with each vial lyophilised proteinase k resuspended in 130ul reconstitution buffer) and incubated at 65C for 6hrs, 75C for 30mins to extract DNA in preparation for sequencing.

### Nanoseq (duplex) sequencing

The lysate from bulk cell sorts (20μl) were submitted to the NanoSeq pipeline for library preparation and sequencing, as has been described in the previous paper^2^. They were then purified using 50μl water and 50μl Ampure XP beads (Beckman Coulter) at room temperature. After a 5-min binding reaction and magnetic bead separation, genomic DNA was washed twice with 75% ethanol, then eluted with 20μl nuclease-free water (NFW). Subsequently, 20 μl of the bead suspension was subjected to an on-bead fragmentation reaction. This fragmentation process took place in a final volume of 25 μl, containing 2.5 μl of 10× CutSmart buffer (consisting of 500 mM potassium acetate, 200 mM Tris-acetate, 100 mM magnesium acetate, 1 mg ml^-1^ BSA, pH 7.9 at 25 °C), 0.5 μl of 5 U μl^-1^ HpyCH4V, and 2 μl NFW. The fragmentation reactions were then incubated at 37 °C for 15 min, followed by purification with 2.5× AMPure XP beads and resuspension in 15 μl NFW. Subsequent A-tailing of the fragmented DNA was conducted in 15 μl reactions, including 10 μl of fragmentation product, 1.5 μl of 10× NEBuffer 4 (consisting of 500 mM potassium acetate, 200 mM Tris-acetate, 100 mM magnesium acetate, 10 mM DTT, pH 7.9 at 25 °C), 0.15 μl of 5 U μl^-1^ Klenow fragment (3′ to 5′ exo-, NEB), either 1.5 μl of 1 mM dATP or 1.5 μl of 1 mM equimolar dATP/ddBTPs, and 1.85 μl NFW. Here, ddBTPs refer to ddTTP, ddCTP, and ddGTP. The reactions were incubated at 37 °C for 30 min. Following this, the 15-μl A-tailing reaction product was combined with 22.4 μl of ligation mix, comprising 2.24 μl of 10× NEBuffer 4, 3.74 μl of 10 mM ATP, 0.33 μl of 15 μM xGen Duplex Seq Adapters (IDT, 1080799), 0.56 μl of 400 U μl^-1^ T4 DNA ligase (NEB), and 15.53 μl NFW. The resulting reactions were incubated at 20 °C for 20 min and subsequently purified with 1× AMPure XP beads, followed by resuspension in 50 μl of NFW.

DNA was quantified by qPCR using a KAPA library quantification kit (KK4835). Samples were diluted in NFW to the standard amount (0.3 fmol for a 15X run) to reach a final volume of 25μl. Subsequently, libraries were amplified using PCR and cleaned up using two consecutive 0.7x AMPure XP beads.

We generated paired-end sequencing reads (150PE) using Illumina NovaSeq platform, resulting in a minimum of 20X per sample.

The sequencing data were then analysed with the BotSeq bioinformatics pipeline (v2.3.2), as described in the previous paper^2^. Sequences were aligned to the human reference genome (hs37d5) using BWA-MEM (v.0.7.5a-r405). Matched normal samples were used to filter out germline SNP from NanoSeq samples. We called indels and normalised the output using bcftools (ref- https://doi.org/10.1093/gigascience/giab008). To assess the authenticity of DNA samples, we used VerifyBamID2^8^, and removed all samples having contamination level > 1% (3 from PD47541, 3 from PD50306, 2 from PD47695). The correction of mutation burden and trinucleotide substitution profiles were generated within the BotSeq pipeline.

### Mutational signature analysis

Hierarchical Dirichlet process (HDP; https://github.com/nicolaroberts/hdp), based on the Bayesian hierarchical Dirichlet process, was used to extract mutational signatures. HDP was run without priors on SBS derived from phylogenetic trees obtained from HSPCs and mutations from NanoSeq samples. The NanoSeq mutations were corrected for the trinucleotide context abundance for each sample. This analysis was performed with two hierarchies, sequencing method and individual id. Both the clustering hyperparameters, alpha and beta, were set to one. The Gibbs sampler was run with 30,000 iterations, spacing of 200 iterations and 100 iterations were collected. After each iteration, three iterations of concentration parameters were performed. Twelve components were extracted, of which five components appeared to be combinations of previously reported signatures. The COSMIC (v3.3) signatures identified were SBS1, SBS5, SBS7a, SBS9 and SBS17. One of the HDP components corresponded to the SBSBlood signature previously reported^9^. Seven components were *de-novo* signatures called predominantly in individuals with chemotherapy. Only signatures with contribution more than 5% of the mutations of the sample’s burden were considered. Signatures with less than 5 percent contribution were termed as overfitting and their contribution was moved under Unassigned.

Eight chemotherapy related signatures derived from HDP (SBSA-SBSH) were examined for their occurrence in exposed vs non-exposed individuals. The proportion of the signatures contributing to the samples in the exposed vs the non-exposed groups were compared using t-test for independent samples, with an equal variance assumption. The test was performed two ways a) per cell type (**Table S6)**; b) combining the samples across cell types for each individual (**Table S7)**. The p-values were adjusted for multiple testing corrections using FDR and Bonferroni methods.

Indel signatures were extracted from small indels called from the HSPC dataset using two different methods. Firstly mSigHdp was run and identified three distinct indel signatures (https://github.com/steverozen/mSigHdp)^10^. To investigate whether the extracted *de novo* signatures were composed of reference COSMIC signatures the SigProfilerAssignment decompose tool was used (https://github.com/AlexandrovLab/SigProfilerAssignment)^11^. Two signatures successfully decomposed, the first into ID1 and ID2 with a reconstructed cosine similarity of 0.99 (compared to the *de novo* signature). The second signature decomposed into ID3, ID5 and ID9 with a reconstructed cosine similarity of 0.93. The final signature, IDA, was initially decomposed into ID2 and ID18 with a reconstructed cosine similarity of 0.88. ID18 is a signature associated with colibactin exposure^12^, which is unlikely in this context. This is in combination with the lower cosine similarity and strong support from the mutation spectra of individuals treated with procarbazine led to this decomposition being rejected.

HDP was also run on the dataset of small indels, without any hierarchy or priors. The hyperparameters and parameters were the same as used for SBS signature extraction. Five ID signatures were extracted, of which one component was similar to COSMIC ID1 signature (ID1; cosine similarity 0.98). Another was a composite of ID3/ID5/ID9 as above. Three novel ID signatures were also extracted: IDA, IDB, IDC; which were associated with platinum, procarbazine and chlorambucil treatments, respectively. Again, only signatures with more than 5% contribution were considered as active. For reporting in the main text we used the more conservative mSigHDP results because of the relatively low numbers of indels in our dataset compared to SBS mutations.

**Supplementary figure 5:**
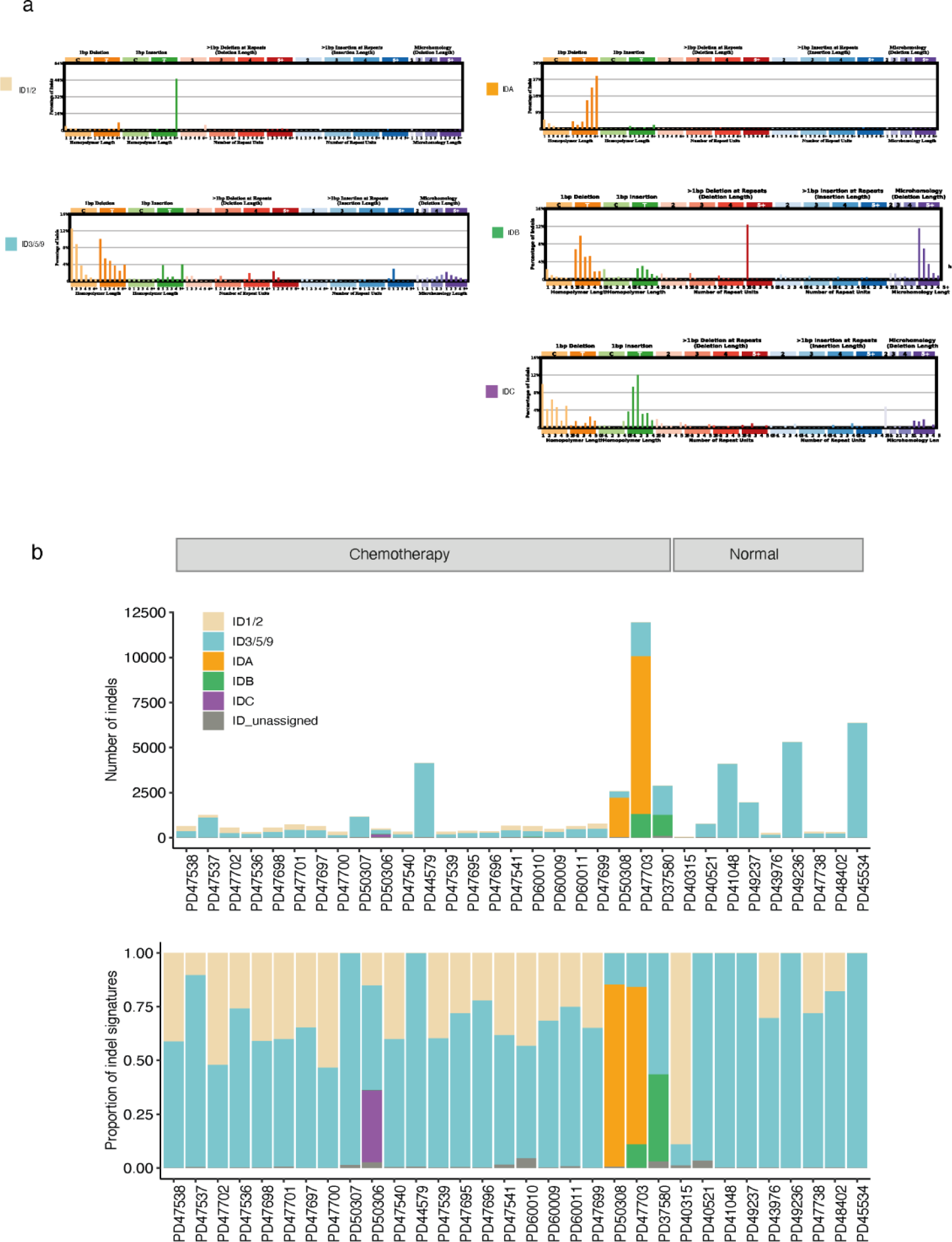
Indel signatures that are present in normal and chemotherapy exposed blood. **a**, Five indel signatures (ID1/2, ID3/5/9, IDA, IDB, IDC) were extracted by HDP. The context on the x-axis show the contributions of different types of indels, grouped by whether variants are deletions or insertions, the size of the event, the presence within repeat units and the sequence content of the indel. **b**, The proportion of indels and indels burden per mutational signatures across 23 chemotherapy exposed and 9 normal individuals, extracting using HDP. Each column represents samples from one individual. Signatures with the contribution <5% are considered as ‘unassigned’.

